# Crossing design shapes patterns of genetic variation in synthetic recombinant populations of *Saccharomyces cerevisiae*

**DOI:** 10.1101/2021.05.26.445861

**Authors:** Mark A. Phillips, Ian C. Kutch, Kaitlin M. McHugh, Savannah K. Taggard, Molly K. Burke

## Abstract

“Synthetic recombinant” populations have emerged as a useful tool for dissecting the genetics of complex traits. They can be used to derive inbred lines for fine QTL mapping, or the populations themselves can be sampled for experimental evolution. In latter application, investigators generally value maximizing genetic variation in constructed populations. This is because in evolution experiments initiated from such populations, adaptation is primarily fueled by standing genetic variation. Despite this reality, little has been done to systematically evaluate how different methods of constructing synthetic populations shape initial patterns of variation. Here we seek to address this issue by comparing outcomes in synthetic recombinant *Saccharomyces cerevisiae* populations created using one of two strategies: pairwise crossing of isogenic strains or simple mixing of strains in equal proportion. We also explore the impact of the varying the number of parental strains. We find that more genetic variation is initially present and maintained when population construction includes a round of pairwise crossing. As perhaps expected, we also observe that increasing the number of parental strains typically increases genetic diversity. In summary, we suggest that when constructing populations for use in evolution experiments, simply mixing founder strains in equal proportion may limit the adaptive potential.

## Introduction

Despite major advances in DNA sequencing technologies, deciphering the genetic basis of complex traits remains a major challenge in modern biology. Efforts to fully characterize the genetic variants underlying traits like human height have had limited success^1^, and there is no clear consensus regarding what models best describe the genetic architecture of complex traits as a whole^2^. Alongside traditional QTL mapping and genome-wide association studies, the “evolve-and-resequence” (E&R) approach has emerged as powerful technique that can generate insight into the genetic basis of complex phenotypes, at least in model organisms^3–5^. This approach involves subjecting experimental populations to conditions that target a specific trait, and simply monitoring the genotypic and phenotypic changes that occur over generations. As these experiments typically feature replicate populations evolving in parallel, researchers can use the data they generate to make powerful statistical associations between genomic variants and traits of interest.

In order to most appropriately model the evolution of complex traits in humans, E&R studies should employ sexually-reproducing eukaryotic organisms^4,6^. However, despite the promise of the E&R approach for dissecting complex traits, there are still unanswered questions regarding what constitutes optimal experimental design. To that end, simulations have assessed the effects of particular experimental parameters on the outcomes of E&R experiments designed to model the evolution of complex traits^7–9^. These simulations support intuitive practices such as maximizing replication and experimental duration, and they recommend maximizing levels of starting genetic variation in experimental populations (i.e. by including more founder haplotypes). Our work here aims to determine how to best achieve this recommendation in experiments with outcrossing *Saccharomyces cerevisiae*.

Most E&R experiments with sexually-reproducing model organisms use *Drosophila*^10–19^. However, outcrossing *S. cerevisiae* populations have emerged as an attractive alternative due to shorter generation times, ease of replication, the ability to freeze and “resurrect” samples taken from different timepoints, and comparable genetic resources^20–21^. As such, we have chosen to focus our efforts on determining how to best construct populations of outcrossing *S. cerevisiae* for use in E&R studies with respect to maximizing standing genetic variation and haplotype diversity.

As *S. cerevisiae* populations isolated from natural or industrial settings typically lack genetic heterogeneity, generating laboratory populations with standing genetic variation requires crossing distinct isogenic strains. These “synthetic recombinant” or multiparent populations have traditionally been used for QTL mapping^22–23^, but they can also serve as the ancestral population in evolution experiments^20–21^. Notably, synthetic recombinant populations have also been used for E&R in other model organisms such as *D. melanogaster* (e.g the *Drosophila* Synthetic Population Resource *cf*. King et al.^24^) and *Caenorhabditis elegans* (e.g. the *C. elegans* Multiparent Experimental Evolution panel, *cf*. Teotónio et al.^25^). In this context, having clearly defined founder genotypes enables the estimation of haplotype frequencies in evolved populations from “Pool-SEQ” data^20,23,26^. And this ability to detect haplotypes responding to selection, as opposed to individual markers, allows for better characterization of linkage and genetic hitchhiking in evolved populations^27^ than is possible in studies where founder genotypes are unknown (e.g. Graves et al.^17^). All of this said, to our knowledge little has been done to explicitly assess how to best generate a recombinant population for downstream use in an evolution experiment. While simulations predict that maximizing the number of founding haplotypes and levels of genetic variation will improve experimental outcomes, there are practical and biological limits to the number of haplotypes that can be combined into a single recombinant background and effectively maintained. One of our goals here is to evaluate the costs and benefits of increasing the number of founding haplotypes in a synthetic recombinant population for E&R.

Another of our goals is to evaluate how different methods of crossing founder strains impact levels of genetic variation in synthetic recombinant populations. There are two general approaches to this task across model organisms. In the more common approach, individuals sampled from various lines are simply mixed in equal proportions and allowed to mate freely prior to the start of the experiment (e.g. Bhargi et al.^19^). With this strategy, the genetic makeup of the resulting population will depend on mating efficiency between individuals from different founding strains, reproductive output, and chance. However, work with *Drosophila* has shown there is no significant allele frequency differentiation between independently constructed synthetic populations using the same sets of isofemale lines^28^. Such a high level of reproducibility could indicate that mating efficiency and differences in reproductive output among founding lines are negligible, or it could indicate that these differences exist and shape the genetic makeup of recombinant populations in a parallel manner. So, while this approach is relatively simple and practical to implement, it may be difficult to maximize the amount of genetic variation and founding haplotype representation in synthetic populations when there are drastic differences in reproductive output and/or mating efficiencies among founding strains. In other words, if there is substantial variation in mating efficiencies among founding strains, this could lead to the over-representation of certain haplotypes belonging to strains that mate most efficiently. By contrast, the second approach for constructing synthetic populations is more complex and involves some level of pairwise crossing between founders^22–25,29^. While this approach is significantly more labor-intensive and time-consuming, it perhaps has advantages in terms of producing populations that have more equal founder haplotype representation and, consequently, higher levels of genetic variation.

Here we assess which of these methods produces populations most well-suited for E&R studies in outcrossing *S. cerevisiae*. We have constructed sets of synthetic populations using both approaches and the same founder populations. In addition to crossing approach, we also evaluate how the number of founder populations used impacts the genetic makeup of the resulting population. Specifically, we created sets of populations using 4, 8 and 12 isogenic strains and both construction methods. Our objective is to characterize patterns of genetic variation and haplotype diversity across these populations to provide recommendations on how to best construct synthetic recombinant populations in outcrossing yeast for the specific application of using them in an E&R study.

## Materials and Methods

### Population creation

All yeast strains used in this study originated from heterothallic, haploid, barcoded derivatives of the SGRP yeast strain collection^30^. A subset of 12 of these haploid strains, originally isolated from distinct geographic locations worldwide, were used to create the synthetic populations we describe here (See Supplementary Fig. S1 for phylogeny). These 12 strains were all genetically modified as described in detail by Linder et al.^23^ to enable easy crossing and diploid recovery; these modified strains were kindly provided by Anthony D. Long (UC Irvine) in 2017. Briefly, these strains were modified so that *MAT*a and *MAT*α strains both contain *ho* deletions to prevent mating-type switching, but each contain a different drug-resistance marker in a pseudogene (YCR043C) tightly linked to the mating type locus (*MAT*a, *ho*Δ, ura3::KANMX-barcode, *ycr043C*::NatMX and *MAT*α, *ho*Δ, ura3::KANMX-barcode, *ycr043C*::HygMX). These genotypes enable haploids of each mating type to be recovered using media supplemented with either hygromycin B or nourseothricin sulfate, and they enable newly mated a/α diploids to be recovered in media supplemented with both drugs.

Two different crossing strategies were used to create genetically diverse populations using 4, 8, and 12 strains as founders (Table 1). “K-type” populations (named for what we call the “kitchen sink method”, or the practice of pooling isogenic strains together without careful focus on representation) were created by simply pooling equal volumes of saturated overnight cultures of the respective haploid founders and allowing those cells to mate. To accomplish this, single colonies of each haploid founder strain were sampled and grown overnight (at 30°C / 200 rpm) in 1 mL of rich media consisting of 1% yeast extract, 2% peptone, and 2% dextrose (YPD). After ~24 hours, cultures were washed in fresh YPD media, pooled with the relevant other overnight cultures in a 50 mL conical tube, vortexed, and now-mixed cultures were allowed to settle and mate for 90 minutes at room temperature. These cultures were then transferred in 200uL aliquots to agar plates containing 100mg/mL nourseothricin sulfate (“NTC”), 300mg/mL hygromycin B (“hyg”) as well as 200mg/mL G418; this strategy ensured that that only newly mated diploids would grow. The resulting lawns of mated diploid cells were collected by scraping with a sterile glass slide into a fresh YPD media. This “cell bank” was archived as a frozen stock at −80°C for each of the K-type populations made with 4, 8, and 12 haploid founders (4K, 8K, and 12K respectively).

**Table 1.** Strains used to create the synthetic populations featured in this study. Strains are arranged in the rows in the table to indicate which specific pairs were crossed in the S-type populations.

| Populations | Founder Strains |  |
| --- | --- | --- |
|  | Mat a | Mat *alpha* |
| S4 & K4 | DBVPG6765 | YPS128 |
|  | DBVPG6044 | Y12 |
| S8 & K8 | DBVPG6765 | YJM981 |
|  | DBVPG6044 | L_1528 |
|  | BC187 | Y12 |
|  | L_1374 | YPS128 |
| S12 & K12 | DBVPG6765 | YJM981 |
|  | DBVPG6044 | Y12 |
|  | BC187 | L_1528 |
|  | SK1 | 273614N |
|  | L_1374 | YPS128 |
|  | YJM975 | UWOPS05_217_3 |

“S-type” populations (named S due to the manipulation of spores to achieve better representation of founder genotypes) were built to mimic more careful crossing designs in which founding lines are crossed in pairs and/or a round-robin. To accomplish this, each haploid strain was paired with a different strain of the opposite mating type and mated as described above. Successful diploid colonies were isolated, grown overnight in 1 mL of YPD, washed and resuspended in sporulation media (1mL 1% potassium acetate), then cultured for 72 hours at 30°C / 200 rpm. Tetrads from these diploid cells were dissected using the Spore Play dissecting microscope (Singer). The four meiotic products (spores) were then collected, allowed to grow for 2 days, and replica plated to plates containing either NTC or hyg to verify the proper segregation of drug resistance markers and thus mating types. Once validated, the meiotic products were grown overnight in 1 mL of YPD. Overnight cultures were standardized to the same optical density (OD_600_) before being pooled in equal volumes in a 50 mL conical tube. Populations were given 90 minutes to mate at room temperature, then were plated on agar plates supplemented with both NTC/hyg/G418 so that only newly mated diploid cells could grow. The resulting lawns of mated diploid cells were collected by scraping with a sterile glass slide into fresh YPD media. This “cell bank” was archived as a frozen stock at −80°C for each of the S-type populations made with 4, 8 and 12 founders (S4, S8, and S12 respectively; see Supplementary Fig. S2 for crossing schematics).

### Population maintenance and 12 cycles of outcrossing

After the creation of the 3 “K-type” and 3 “S-type” synthetic recombinant populations described above, all populations were taken through 12 consecutive cycles of intentional outcrossing; in other words, the populations were subjected in parallel to a series of steps that induced regular sporulation, spore isolation, and mating. Detailed methods are described by Burke et al.^31^. Briefly, newly mated diploid cells from the last step of the population creation protocol (i.e. the “cell banks”) were grown overnight in 10 mL YPD media. These cultures were washed and resuspended in sporulation media, and incubated with shaking for 72 hours (30°C/200 rpm). Cells then underwent a number of methods to disrupt asci and isolate/randomize spores, including incubation with Y-PER yeast protein extraction reagent (Thermo) to kill vegetative diploids, digestion with 1% zymolyase (Zymo Research) to weaken ascus walls, and well as high-speed shaking with 0.5 mm silica beads (BioSpec) to mechanically agitate the asci. After these steps spores were resuspended in 10 mL YPD and allowed to settle and mate for 90 minutes at room temperature. Diploids were recovered as described above; cultures were transferred to 10 individual YPD agar plates supplemented with NTC/hyg/G418 in 200 μL aliquots and incubated at 30°C for 48 hours. The resulting lawns of mated diploid cells were collected by scraping with a sterile glass slide into fresh YPD media. This “cell bank” was again sampled for archiving at −80°C, and used to initiate an overnight culture for the next outcrossing cycle. We estimate that 15–20 asexual generations occurred between every outcrossing cycle of the experiment. Based on counting colonies from dilutions of cultures plated at various benchmarks during the protocol, we expect that 7.5–11 generations elapse during the overnight culture in YPD media, and another 7.5-11 generations elapse during the period of diploid recovery on agar plates. Thus, a minimum of 15*12 = 180 cell doublings likely took place over the 12 cycles of outcrossing in each synthetic recombinant population.

### Genome sequencing and SNP identification

Each of the recombinant K- or S-type population was sequenced at three specific timepoints: initially (we also call this timepoint “cycle 0”), after 6 cycles of outcrossing (“cycle 6”), and after 12 cycles of outcrossing (“cycle 12”). We also sequenced each haploid founder strain such that we could estimate the relative contributions of each to the recombinant populations. Each of the founding SGRP strains were plated as haploids on plates containing either NTC or hygromycin to verify the presence of the appropriate drug resistance markers. Individual colonies were isolated from each strain for verification of identifying barcodes at the URA3 locus using Sanger sequencing (Cubillos et al.^30^ provide barcode and primer sequences). Once validated, single colonies were again isolated for whole-genome sequencing. One milliliter of YPD media was inoculated with single colonies, grown overnight, and the resulting culture was harvested for gDNA extracted using the Qiagen Puregene Yeast/Bact. Kit. Purified gDNA from each haploid founder was then prepared for sequencing using the Nextera DNA Sample Preparation Kit (Illumina). Some minor modifications to the manufacturer’s protocol were implemented to optimize throughput (*cf*. Baym et al.^32^). Genomic DNA libraries were prepared for experimental recombinant populations in the same way and all samples were pooled to generate a single multiplexed library. Because the recombinant (i.e. genetically variable) populations require significantly higher coverage to accurately estimate allele frequencies at variable sites, these populations were added to the library at 10X the molarity of each haploid founder sample. The multiplexed library was run on two SE150 lanes on the HiSeq3000 at the OSU Center for Genomic Research and Biocomputing (CGRB). Data for the 4S populations were previously published in Burke et al.^31^ and raw fastq files are available through NCBI SRA (BioProject ID: PRJNA678990). Raw fastqs for all other populations are also available through NCBI SRA (BioPorject ID: PRJNA732717).

We have developed a processing pipeline for estimating allele frequencies in each population directly from our pooled sequence data. We used GATK v4.0^33–34^ to align raw data to the *S. cerevisiae* S288C reference genome (R64-2-1) and create a single VCF file for all variants identified across all replicate populations, using standard best practices workflows and default filtering options. We also downloaded and indexed a reference VCF file with SNP information for a number of distinct natural isolates of S. cerevisiae^35^; this is a recommended best practice for calibrating base quality with GATK v4.0. This VCF file was converted into a SNP frequency table by extracting the AD (allele depth) and DP (unfiltered depth) fields for all SNPs passing quality filters; the former field was used as the minor allele count and the latter was used as the total coverage. The python scripts used to generate and convert VCF files to tables suitable for downstream analyses in R (www.R-project.org) are available through GitHub (see Data Availability statement for details on where to find all major scripts used to process and analyze data).

Our general SNP analysis strategy involved portioning the data to create three separate SNP tables with each table corresponding to a set of founders and populations derived from them (e.g. a table containing with the S4 and K4 populations and their founders). In each table, we chose to only include sites with a minimum coverage >20X in the in synthetic populations as a quality control measure. Next, sites were filtered based on data from the founder populations. We excluded all sites that appeared to be polymorphic within a given founder, and sites where a single nucleotide was fixed across all founders. This was done as such occurrences could indicate sequencing error given that our founder strains are haploid and isogenic, and a site is unlikely to be polymorphic in our synthetic populations if it is fixed across all of the founders. After these filters were applied, we retained a collection of high-quality SNPs in each population to subject to further analysis. The total number of SNPs identified in each population is given in Table 1, and the average genome-wide coverage (i.e. depth of sequence coverage) of each population is given in Supplementary Table S1. All populations had mean coverages >50X with all but one population (S4 cycle 0) having greater than 70X mean coverage (Supplementary Table S1).

### SNP Variation

Our main objective was to evaluate how crossing strategy and the number of founder strains impacts patterns of SNP variation in synthetic recombinant populations. To that end, we assessed SNP-level variation in our recombinant populations using several metrics. First, we simply determined the number of polymorphic sites segregating in each population immediately following their creation (cycle 0), and monitored how that number changed over time ((i.e. after 6 or 12 outcrossing cycles). This approach of tracking the total number of SNPs should reveal whether particular crossing strategies – i.e. using a certain number of founders, and/or one of the two crossing strategies – consistently produced populations with more SNPs, and whether these SNPs were maintained or lost over 12 outcrossing cycles. We also generated UpSet plots using the UpsetR package^36^ in R to visualize patterns of overlap between the total number of SNPs possible for a given combination of founder strains, and the SNPs we observed in our actual populations. We define the total number of possible SNPs as all loci for which at least one of the founding strains used has an allele different from the others; this number will therefore differ among the 4-way, 8-way, and 12-way crosses.

In addition to SNP number, we also characterized the distribution of SNP frequencies in each population, which allows more direct comparisons between populations with different numbers of founders but the same crossing strategy, or the same number of founders but different crossing strategy. To do this, we focused on two metrics: the site frequency spectrum (SFS), and genome-wide heterozygosity. Here heterozygosity refers to *2pq*, the product of the reference (i.e. the S288C allele) and alternate allele frequency at a given site multiplied by 2. In addition to looking at differences in mean genome-wide heterozygosity between populations, we also generated sliding window plots showing patterns of variation across each chromosome. To define windows, we used the GenWin package^37^ in R with the following parameters: “smoothness = 6000, method = 3.” GenWin itself uses a smoothing spline technique to define windows based on breakpoints in the data. While we ultimately used “smoothness = 6000”, we did initially try a range of values. Our final selection was made based on what most clearly represented trends in the data. For interested parties, plots with more or less smoothness can be easily generated using data and scripts we have made available through Dryad and Github (See “Data availability” statement for details).

It is worth noting that our ability to assess levels of genetic variation across our synthetic populations is limited by the fact we have only collected Pool-SEQ data. Given the complex life-history of the yeast populations in this experiment, which involves periods of 7-15 generations of asexual growth punctuated by discrete outcrossing events, it is not possible for the genotypes of all individuals in the population to be shuffled by recombination every generation. Therefore, asexual lineages will evolve by clonal interference for relatively short periods of time, until the next outcrossing event decouples individual adaptive alleles from a particular genetic background. It is possible that during these periods of clonal interference, particular diploid lineages will dominate, and if these lineages are heterozygous at a given locus, that will lead to an artificially elevated heterozygosity value at that SNP. But we do not believe this is a major complication in the understanding of nucleotide diversity in our experiment for several reasons; namely, that our outcrossing protocol includes several measures that maximize outcrossing efficiency (i.e. any asexual diploids that fail to sporulate are killed), the generally high rate of recombination in yeast, and that the periods of asexual growth are short and unlikely to exceed ~20 cell doublings.

### SNP frequency changes over 12 cycles of outcrossing

Although statistical power in this power is limited due to a lack of replication, we attempted to identify regions of the genome showing obvious responses to selection in each synthetic population. Specifically, we used Pearson’s χ^2^ test as implemented in the poolSeq^38^ package in R to compare SNP frequencies between cycle 0 and cycle 12 in each synthetic population. We chose this particular test based on a benchmarking effort that suggests it is well-suited to detecting selection in E&R experiments lacking replication^9^. After results were generated for each synthetic population, log transformed p-values were plotted for each chromosome across sliding windows. The GenWin package in R (parameters: “smoothness = 2000, method = 3”) was once again used to define windows based on breakpoints in the data. Plots were then examined to see if there were any genomic regions showing signs of selection based on significance levels relative to the background.

### Haplotype representation

In addition to describing SNP diversity, we also describe the diversity of founder haplotypes represented in our synthetic populations. We were particularly interested in evaluating whether the S-type strategy might produce populations in which founder haplotypes are more evenly represented (at intermediate frequency) compared to the K-strategy. Given the stochasticity inherent in the K-type strategy, we thought it probable that founder genotypes with especially high sporulation and/or mating efficiencies (i.e. those with the highest reproductive outputs in the outcrossing context) might come to dominate. To this end, we estimated haplotype frequencies in all experimental populations initially, and after 6 and 12 cycles of outcrossing to determine how evenly haplotypes were represented, and how this might have changed over time. We used the sliding-window haplotype caller described in Linder et al. (2020) and software the authors have made available as a community resource: https://github.com/tdlong/yeast_SNP-HAP. Our results were generated by using the haplotyper.limSolve.code.R script and estimates were made across 30KB windows with a 1KB stepsize. This particular haplotype caller was developed specifically to estimate haplotype frequencies in multiparent populations when founder haplotypes are known. A full description of the algorithm being used, and results of empirical validation can be found in Linder et al. (2020). To quantify haplotype variation in each population, we calculated haplotype diversity (H) using the following formula:

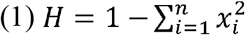

where *x_i_* is the frequency of the *i*th haplotype of the *n* founders used to create given population^39^. Though it is worth noting that maximum expected H will vary depending on the number of founders used to create a given population as

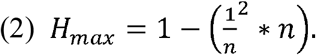

### Phenotypic characterization of experimental populations

To evaluate the possibility that populations might be phenotypically differentiated, we measured two life-history traits: sporulation efficiency and growth rate._We estimated the 3-day sporulation efficiency for each recombinant population at the beginning and end of the experiment, as this is a life-history trait that might have reasonably responded to the selection imposed by the regular outcrossing protocol. All populations archived at “cycle 0” (i.e. the pool of diploid cells used to initiate each K- or S-type population) and “cycle 12” (i.e. diploid cells recovered from each population after the 12^th^ outcrossing cycle) were revived by plating 1mL of thawed culture onto a YPD agar plate and incubation at 30°C for 48 hours. In order to sample the genetic diversity of each population, a sterile wooden applicator was scraped in a zig-zag pattern across the lawn of cells on each plate to collect a pinhead-sized clump of yeast. Each clump was mixed in 10 mL YPD in a 50 mL conical tube and vortexed. Tubes were then incubated at 30°C/200 rpm for ~24 hours. After confirming that each tube had comparable cell densities – this was done by verifying that the OD_600_ absorbance value of a 1:100 dilution ranged between 0.095-0.2 – cell pellets were collected by spinning for 5 minutes at 5000 rpm. Cell pellets were washed in 1 mL of sterile water, spun down again, and resuspended in 40 mL of minimal sporulation media (1% potassium acetate w/v). Each culture was transferred to sterile 250 mL Erlenmeyer flasks and covered loosely with foil, where they were cultured at 30°C/200 rpm for ~72 hours to sporulate. After sporulation, aliquots of each culture were loaded onto a hemacytometer (Incyto C-Chip, type NI) and visualized under 40x magnification on a Singer SporePlay microscope. For each culture, ~200 cells were counted (specific range: 190-230 cells), and sporulation efficiencies were estimated as the proportion of tetrads observed over the total number of cells in the field of view. Sporulation efficiency for each of the 12 recombinant populations (6 “cycle 0” and 6 “cycle 12”) was assessed by averaging these proportions over 2-3 independent biological replicates.

In addition to characterizing sporulation efficiencies for each of the “cycle 0” and “cycle 12” recombinant populations, we also measured growth rate with high-throughput absorbance-based assays in liquid YPD. We also included the 12 founder strains in this assay, for comparison with the recombinant populations. S- and K-type recombinant populations were sampled from each freezer recovery plate as described above. Haploid founder strains were revived from freezer stocks by striking for single colonies onto YPD agar plates. Each population or strain was assayed in two biological replicates; recombinant populations were sampled to inoculate two separate overnight cultures in liquid YPD, and strains were sampled by picking two distinct colonies to initiate two separate overnight cultures (one colony per culture). All biological replicates were incubated for ~24 hours at 30°C/200 rpm. The day of the assay, OD_600_ was measured in all cultures and the readings used to standardize them to a target OD_600_ of 0.05 in fresh YPD (observed values ranged 0.042-0.061). 200uL of each culture was aliquoted to separate wells of a 96-well plate, with two technical replicates per biological replicate. The arrangement of technical replicates on the plate was carried out in an attempt to control for possible edge effects. The growth rate assay was carried out in a Tecan Spark Multimode Microplate Reader, set to record the absorbance at 600 nm for each well every 30 minutes for 48 hours at 30°C, without plate agitation/aeration. The R-package “Growthcurver” (Sprouffske and Wagner^40^) was used to estimate population growth parameters from the raw data. In order to determine the carrying capacity and doubling time of the culture in each well, the absorbance measurements taken during the assay were fit to the following equation:

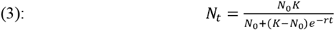

Where *N_t_* is the absorbance reading at time *t*, *N*_0_ is the initial absorbance, *K* is the carrying capacity, and *r* is the growth rate, or doubling time. Here, doubling time refers to the time necessary for the size of a population to double under non-restricted conditions, while carrying capacity is the maximum population size under the given conditions. The values for each biological replicate were averaged across technical replicates, and the values for each strain/population were determined by averaging across biological replicates.

## Results

### SNP Variation

To assess how crossing strategy and number of founder strains impacts SNP variation, we began by simply counting the number of SNPs present in each of our synthetic populations upon their creation and how that changes over several cycles of recombination (Table 2). As expected, the total number of possible SNPs that can possibly contribute to segregating genetic variation increases with the number of founders used. Looking at our actual populations at cycle 0 and focusing on those created using the same crossing strategy, we also generally find the observed number of SNPs in each population to increase with the number of founders used. The only exception to this pattern is the K12 population where we see dramatic losses in polymorphic sites relative to all other populations. We also typically observe reductions in the number of SNPs in all experimental populations over time. However, we do note higher “stability” (i.e. smaller losses) in the 8-founder populations, and in population S8, we actually observe higher SNP counts in cycle 12 than in cycle 6. This discrepancy is most likely due to a relatively small number of sites at very low frequency in cycle 6 (i.e. too low for our SNP calling to pick up), increasing to detectable levels by cycle 12. Nevertheless, the overall trend still appears to be reductions in the number of polymorphic sites over time. Our data also suggest these reductions are typically more pronounced in populations created using the K-type strategy, and that populations created using the S-type strategy have more polymorphic sites than those created with the K-type strategy.

**Table 2.** Number of possible SNPs and those actually observed in all synthetic populations at each cycle where samples were taken for sequencing. Percentages are relative to total possible number of SNPs for a given set of founders (n=4, 8, or 12). We estimate that approximately 15-20 asexual generations occur between each cycle of outcrossing.

|  | **S4** | **K4** | **S8** | **K8** | **S12** | **K12** |
| --- | --- | --- | --- | --- | --- | --- |
| **Possible SNPs** | 112,738 |  | 122,663 |  | 155,516 |  |
| **Cycle 0** | 91,658<br>(81%) | 71,771<br>(64%) | 98,864<br>(81%) | 94,965<br>(77%) | 101,134<br>(65%) | 50,352<br>(32%) |
| **Cycle 6** | 75,712<br>(67%) | 69,954<br>(62%) | 78,078<br>(64%) | 62,537<br>(51%) | 79,070<br>(51%) | 38,853<br>(25%) |
| **Cycle 12** | 57,405<br>(51%) | 39,085<br>(35%) | 80,941<br>(66%) | 61,636<br>(50%) | 73,894<br>(48%) | 5,171<br>(3%) |

We also examined patterns of overlap between our synthetic populations and what is possible given their respective founders using UpSet plots (Supplementary Fig. S3). UpSet plots are useful because they allow visualization of observations unique to groups; in other words, they highlight observations that are included in a specific group and excluded from all others. As an illustrative example, the fourth vertical bar in Supplementary Fig. S3A-C represents all of the SNPs that could possibly segregate in a population but do not, and this reveals that a greater proportion of possible SNPs are lost in synthetic populations with 12 founders compared to those with 4 or 8 (this result is also noted in Table 2). We were interested in using this visualization approach to evaluate whether there might be particular SNP groups that are hallmarks of the S-type or K-type strategy. While many more of the possible SNPs appear in S-type populations relative to the K-type populations for a given set of founders, the UpSet plots indicate that very few SNPS are unique to a particular crossing strategy. We interpret this result as evidence that neither crossing strategy favors specific alleles, and this is true regardless of the number of founder haplotypes used.

Next, we looked at how the SFS varied across different populations and how they changed over time. As shown in Fig. 1, the S-type populations tend to exhibit less skewed frequency distributions compared to the K-type population, and they are also more stable over time. This is particularly evident in the 12-founder populations. In these we see that by the final cycle of recombination in the K12 populations, there is an extreme skew in the SFS with most sites exhibiting very high or low SNP frequencies. This is also consistent with the results shown in Table 2 where we see much higher levels of fixation over time in K12 than in any other population. This contrast is present but far less extreme in populations created with 4 or 8 founder strains. In general, at cycle 0 all populations deviate substantially from the SFS that we would expect if the respective founders combined in perfectly equal proportions (Supplementary Fig. S4). These deviations suggest that drift and/or selection are impacting the genetic makeup of synthetic populations from the moment they are established.

**Figure 1.**
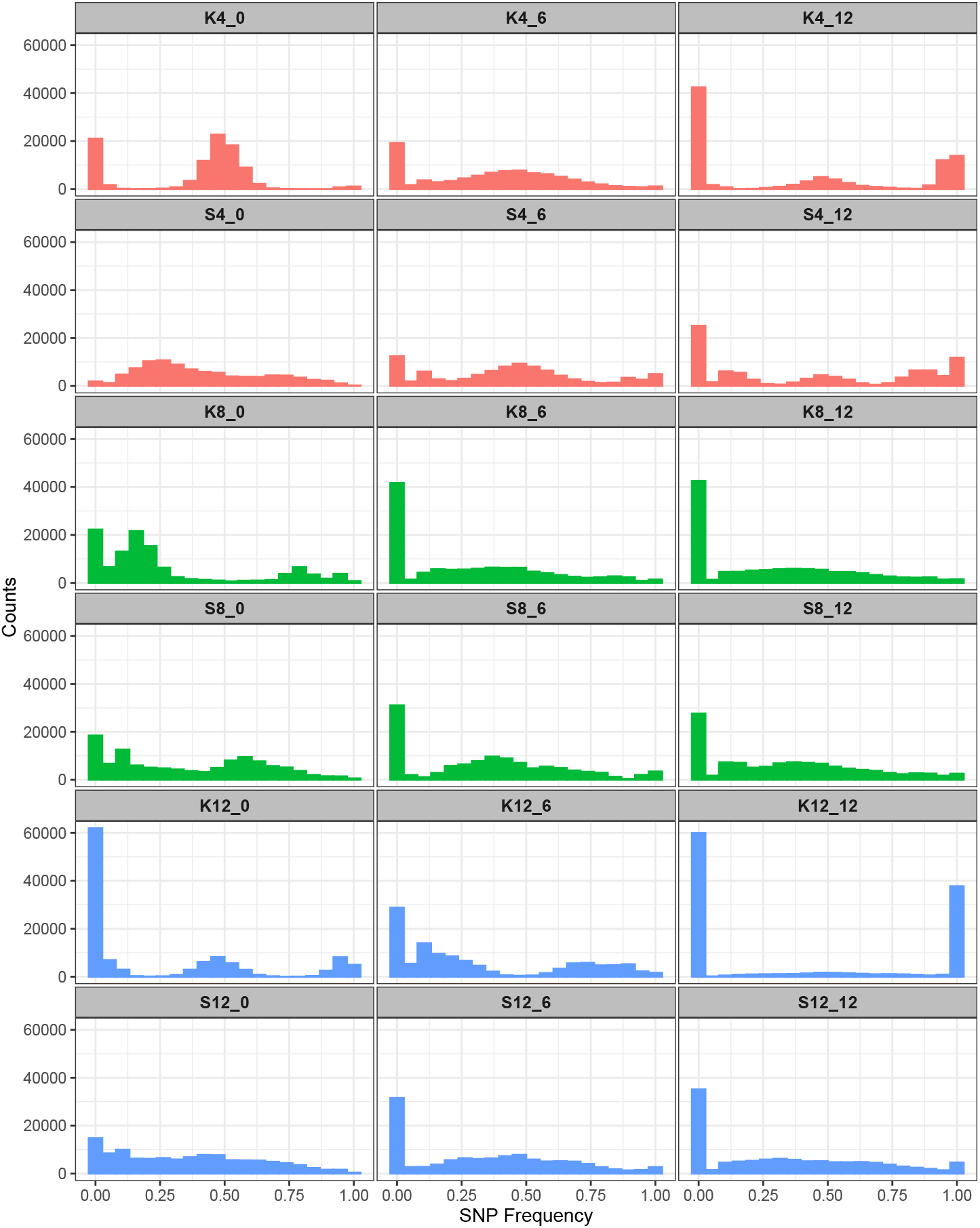
Site frequency spectra (SFS) for all populations at each timepoint samples were taken for DNA sequencing; immediately after construction, or “cycle 0” (left panels), after 6 cycles of outcrossing (middle panels), and after 12 cycles of outcrossing (right panels).

Finally, we assessed the effects of crossing strategy and number of founders on genome-wide heterozygosity. The clearest pattern we observed is that after 12 cycles of recombination, S-type populations exhibited greater overall levels of heterozygosity across the genome compared to their K-type counterparts (Table 3). After 12 cycles of outcrossing, we also see large stretches along chromosomes where heterozygosity is near zero in K4 and K12 which is not the case in their S-type counterparts (Fig. 2). Other patterns in the data are less clear, however; for instance, we do not find stretches of the genome where variation has been expunged in the K8 population, compared to the S8 counterpart (Fig 2C-D). Looking at the S-type populations alone, we find that by cycle 12 the S4 populations have experienced a greater loss of heterozygosity than the S8 and S12 populations (Table 3). However, differences between S8 and S12 populations are far less severe with the former having slightly higher mean heterozygosity. As such, there is no clear positive relationship between heterozygosity and number of founders. This pattern largely breaks down in the K populations. The K4 population experiences a greater loss of heterozygosity than K8 by cycle 12, but then K12 experiences the most severe declines in heterozygosity by cycle 12 as expected given the other measures of SNP variation we have looked at thus far.

**Table 3.** Mean genome-wide heterozygosity for all synthetic populations at each cycle where samples were taken for sequencing. Mean heterozygosity is often, but not always, higher in the S-type populations (shaded gray) relative to their K-type counterparts at a given timepoint.

|  | **S4** | **K4** | **S8** | **K8** | **S12** | **K12** |
| --- | --- | --- | --- | --- | --- | --- |
| **Cycle 0** | 0.38 | 0.47 | 0.36 | 0.28 | 0.36 | 0.31 |
| **Cycle 6** | 0.32 | 0.41 | 0.33 | 0.25 | 0.32 | 0.25 |
| **Cycle 12** | 0.20 | 0.17 | 0.31 | 0.25 | 0.28 | 0.04 |

**Figure 2.**
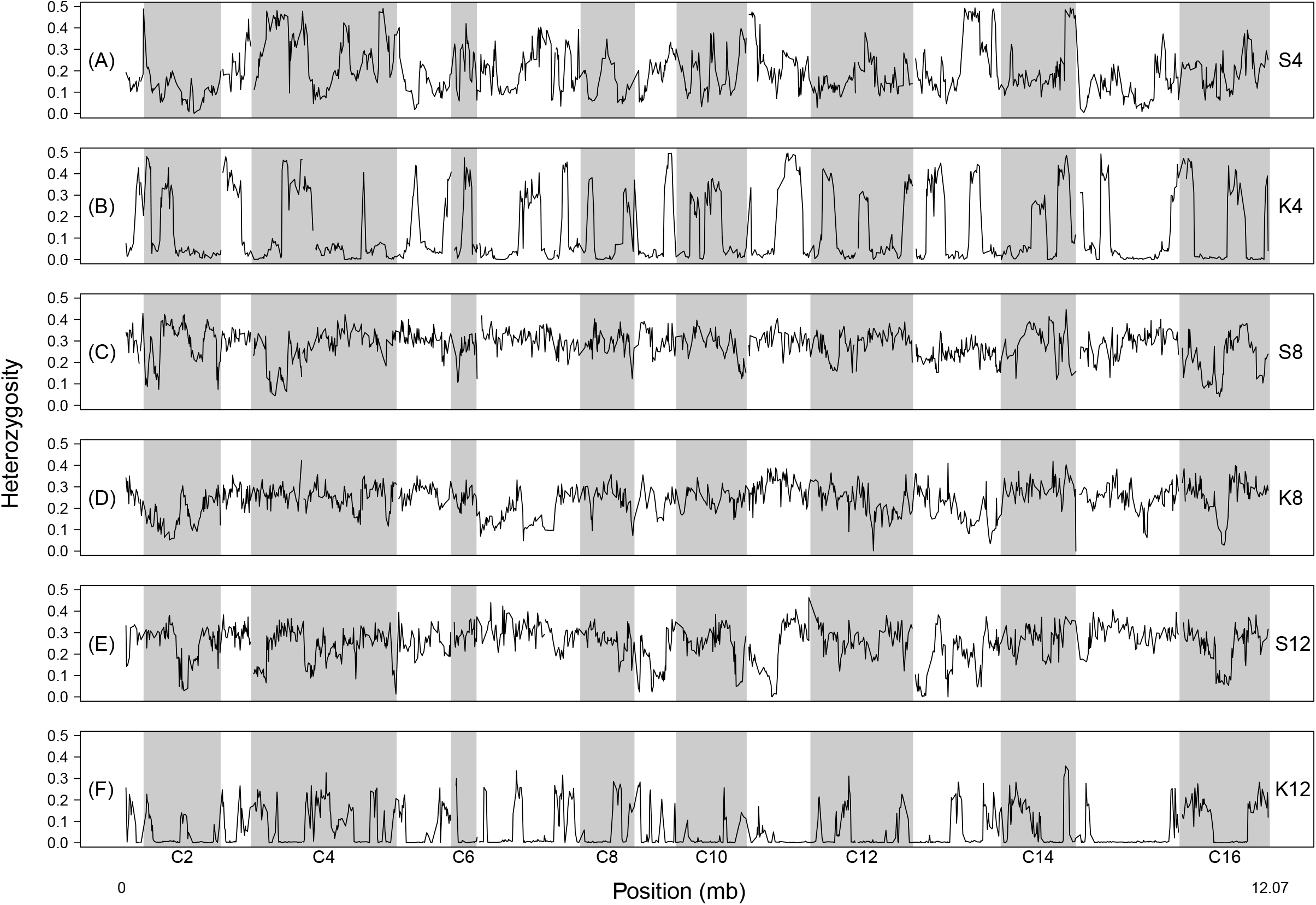
Sliding window heterozygosity for all populations featured in this study after 12 cycles of outcrossing.

### SNP frequency changes over 12 cycles of outcrossing

We used Pearson’s χ^2^ test to compare SNP frequencies between cycle 0 and cycle 12 of each individual population to see if there were any regions of the genome showing obvious responses to selection imposed by our outcrossing maintenance protocol. Looking across results for each population, we do not find any genomic regions that show consistent responses to selection (Fig. 3). However, we do find instances in individual populations where there are clear peaks in significance relative to the rest of the genome (e.g. Fig. 3A and C-E). Widespread fixation in K4 and K12 make it difficult to identify such peaks (Fig. 3B and F).

**Figure 3.**
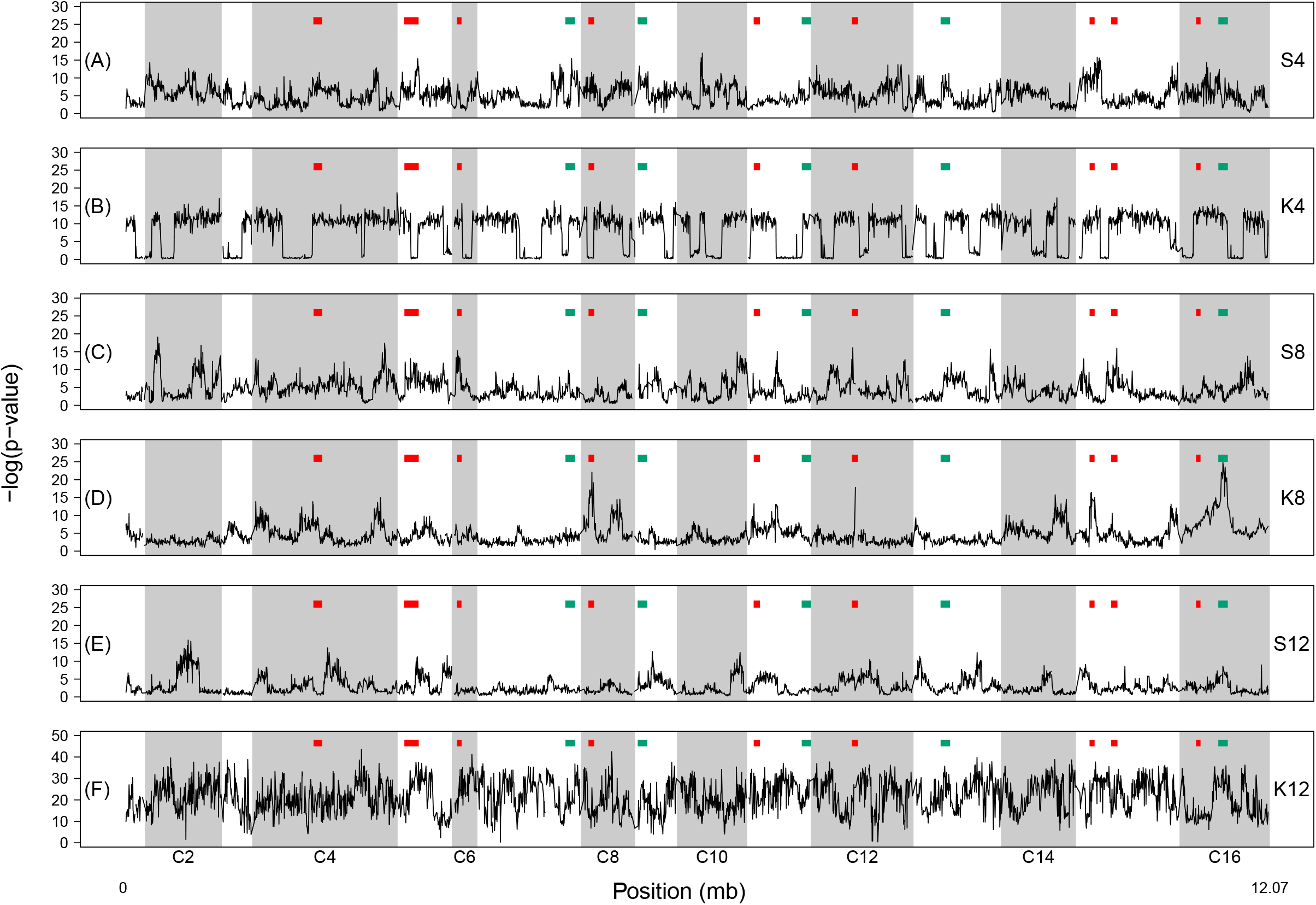
Results from Pearson’s χ^2^ test comparing SNP frequencies initially (cycle 0), and after 12 cycles of outcrossing for all populations featured in this study. In each panel, red and green boxes indicate regions of genome and genes associated with selection for frequent outcrossing from Cubillos et al. (2013) and Burke et al. (2014), respectively

We also compared our results to prior work which identified regions of the genome and genes associated with selection for frequent outcrossing, Cubillos et al.^22^ and Burke et al.^20^. Our rationale for doing this is that as our current study features no within-treatment replication, it is weak to implicate signatures of adaptation by itself; but, obvious overlap with other studies would provide indirect evidence for adaptation. Similarly, if a particular population (or population type) implicated more candidate regions from the literature than the others, this could provide evidence that a particular crossing strategy might lead to the best outcomes in an E&R study, with respect to identifying regions of interest. We do find instances where peaks in significance align with previously identified candidate regions (e.g. Fig. 3A, C-D), but there is no clear pattern where peaks consistently line up with a given candidate region across multiple populations. There are also many more instances where we do not find peaks that align with the previously identified candidate regions. So generally speaking, while this comparative approach provides opportunities for describing particular regions, perhaps those of *a priori* interest, we cannot conclude that observed changes in SNP frequencies in any population are likely signatures of selection for forced outcrossing. On the other hand, we also observe that the 8-founder populations generally implicate more candidate genes from the literature, compared to any other population (Fig. 3C-D). This suggests that perhaps using 8 founding haplotypes in a population for an E&R project leads to better outcomes, in terms of identifying candidate regions, than using 4 or 12. Notably, the K8 population and S8 population implicated similar numbers of peaks, so it is not clear that one crossing strategy is better than the other in this respect.

### Haplotype Representation

Using sequence data from all experimental populations and our founder strains, we estimated haplotype frequencies across the genome to assess how crossing strategy and number of founders impact haplotype representation initially, as well as after 6 or 12 cycles of outcrossing. Notably, estimates for K12 are made using far fewer SNPs due to the extreme levels of fixation seen in this population and are almost certainly less reliable than those from other populations. We first assessed the haplotype frequencies observed after 12 cycles of outcrossing, as this provides a birds-eye view of the amount of haplotype diversity present at the end of the experiment (Supplementary Figs. S5-S10). While haplotype frequencies fluctuate across the genome in all populations, mean genome-wide haplotype frequency estimates point to clearer patterns. In the 4- and 12-founder populations, we find that the S-type populations have haplotype frequencies that are more evenly-distributed compared to the K-type populations (Table 4). This is also reflected in levels of haplotype diversity for these populations, where we find that the S-type populations typically have greater levels of diversity across the genome compared to the K-type populations (Fig. 4). Mean haplotype diversity is typically greater in the S-type populations, closer to maximum expected values, and has smaller variance across the genome (Supplementary Table S2). These differences can largely be attributed to the almost complete loss of particular haplotypes in the K-type populations versus their S-type counterparts. For instance, in 4S we observe nearly equal representation of the founding haplotypes (Table 4; Supplementary Fig. S5), but in K4 we observe that two of the founding haplotypes dominate and the other two are almost entirely lost (Table 4; Supplementary Fig. S6). The S8 and K8 populations appear more similar in measures of haplotype diversity (Fig. 4; Supplementary Table S2, but as with K4, we once again that two haplotypes, DBVPG6044 and YPS128, are almost entirely lost (Supplementary Fig. S8; Table 4) in a way not seen in S8 (Supplementary Fig. S7; Table 4). The S12 (Supplementary Fig. S9) and K12 (Supplementary Fig. S10) results show similar patterns, but interpretation is complicated by the fact K12 haplotype estimates were generated using a very limited number of SNPs. However, across our study as a whole, haplotype representation does appear to benefit from an S-type crossing strategy. Using fewer founders as well as an S-type crossing strategy yields distributions that most closely match what we would expect under equal blending.

**Table 4.**
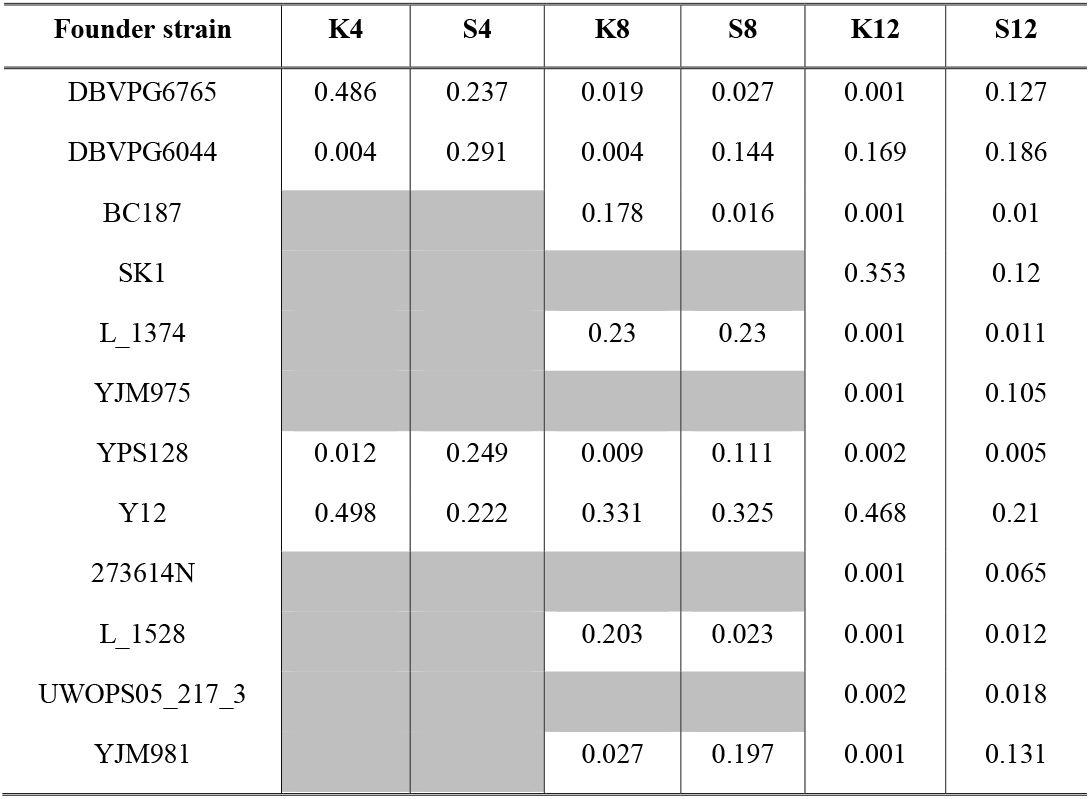
Mean genome-wide haplotype frequencies for all founder populations in each synthetic population based on SNP frequencies observed after 12 cycles of outcrossing (*note*: shaded cells indicate that a founder not used for that particular population).

| Founder strain | K4 | S4 | K8 | S8 | K12 | S12 |
| --- | --- | --- | --- | --- | --- | --- |
| DBVPG6765 | 0.486 | 0.237 | 0.019 | 0.027 | 0.001 | 0.127 |
| DBVPG6044 | 0.004 | 0.291 | 0.004 | 0.144 | 0.169 | 0.186 |
| BC187 |  |  | 0.178 | 0.016 | 0.001 | 0.01 |
| SK1 |  |  |  |  | 0.353 | 0.12 |
| L_1374 |  |  | 0.23 | 0.23 | 0.001 | 0.011 |
| YJM975 |  |  |  |  | 0.001 | 0.105 |
| YPS128 | 0.012 | 0.249 | 0.009 | 0.111 | 0.002 | 0.005 |
| Y12 | 0.498 | 0.222 | 0.331 | 0.325 | 0.468 | 0.21 |
| 273614N |  |  |  |  | 0.001 | 0.065 |
| L_1528 |  |  | 0.203 | 0.023 | 0.001 | 0.012 |
| UWOPS05_217_3 |  |  |  |  | 0.002 | 0.018 |
| YJM981 |  |  | 0.027 | 0.197 | 0.001 | 0.131 |

**Figure 4.**
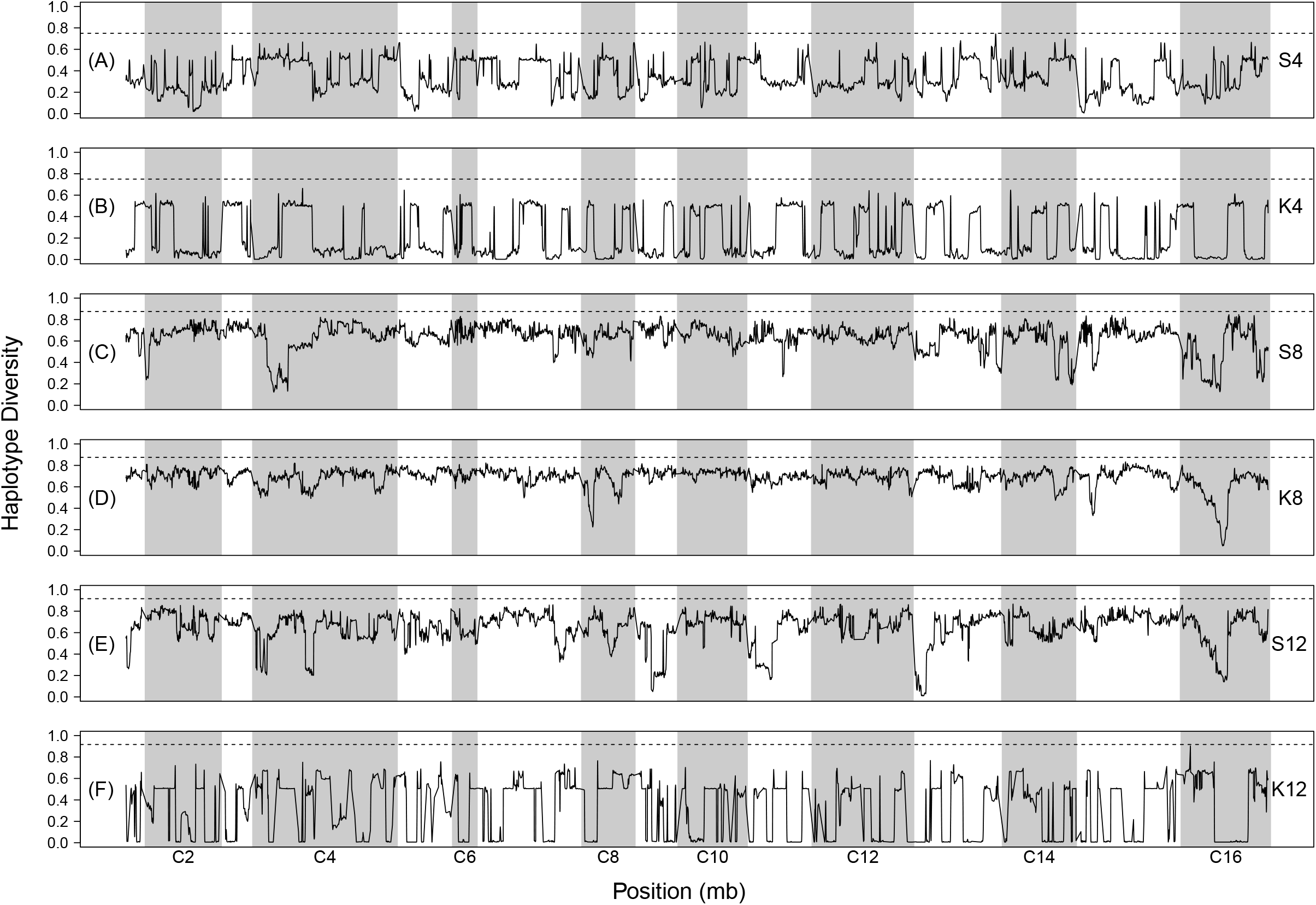
Haplotype diversity for all populations featured in this study after 12 cycles of outcrossing. The dotted line in each panel indicates the maximum expected haplotype diversity for each population.

While patterns of haplotype representation in the “cycle 12” populations speak to how well haplotype diversity is maintained over time, evaluating haplotype representation at earlier parts of the experiment also reveals insight into how the K- versus S- strategies impact haplotype diversity. The near-complete absence of the YPS128 and DBVPG6044 haplotypes in K4 after 12 outcrossing cycles begs the question of exactly when these haplotypes were lost. In fact, these two haplotypes were not observed at high frequencies even immediately after the population’s founding, which suggests that these two strains simply failed to mate with the other strains in the pool (Supplementary Figure S11). The general pattern of haplotype frequencies in this population over time suggests that these two haplotypes gradually diverge from starting frequencies near ~0.5 at cycle 0 (Supplementary Figure S11), such that at cycle 6 frequencies are more variable (Supplementary Figure S12) and by cycle 12 the frequencies are all nearly fixed at any given position along the genome. This is in stark contrast to what we see in S4 where all founders are well-represented at each timepoint (Supplementary Figure S7, S13-14). However, patterns become more complicated in populations created using more founders. For instance, the haplotype frequencies in the initial K8 population (Supplementary Figure S15) suggest that YPS128 was lost immediately, as we also do not observe it after 6 cycles (Supplementary Figure S16) or 12 cycles (Supplementary Figure S8) of outcrossing. But Y12, which is also appears lost in the initial population, increases after 6 and 12 cycles to become one of the most prevalent haplotypes in the population. We also see the opposite pattern for DBVPG6044. The observation that a single founder is lost for good in one K-type population but recovers in another is difficult to explain, but it is consistent with the idea that the K-type strategy is generally more unpredictable, and perhaps more prone to haplotype loss than the S-type strategy.

Next, we evaluated whether any of our observed patterns of haplotype representation could be easily explained by selection. To do this we compared our findings to the results of Burke et al.^20^. In this study, twelve replicate populations were created using the same four founding strains as the S4 and K4 populations, and were subjected to a similar outcrossing regime for 18 cycles (Note: the synthetic population the replicates were derived from was created using a S-type strategy and underwent 12 cycles of recombination before replicates were generated). Based on their analysis of genomic data taken from each population across three timepoints, Burke et al. (2014) identified five major candidate regions associated with adaptation to forced outcrossing. Four of these regions (regions A, B, C, and E in the paper) were also found to be clearly defined by a given haplotype at the end of the experiment (See Supplementary Figure S4 in Burke et al.^20^). As the founders used to create the synthetic populations of Burke et al. (2014) are present in all of the populations featured in this study, we can reasonably interpret changes in haplotype frequency in these four candidate regions in our populations as evidence of selection. In other words, as our populations here are unreplicated, and we cannot use strong-inference approaches with them directly, we sought parallels between haplotype frequency differences observed here and those described as adaptive in a very similar experiment.

As shown in Table 5, we find some evidence that the haplotypes driving adaptation in the Burke et al.^20^ study also show substantial increases in frequency in our populations. In candidate regions A (C9: 950,000-975,000) and E (C13:445,000-460,000), we find clear evidence of the Y12 haplotype frequency increasing across all of our populations as it did in the Burke et al. (2014) study. Given how dramatic some of these changes are from cycle 0 to cycle 12, we think it reasonable to speculate that these haplotypes might also be driving adaptation here. However, this pattern in not corroborated for candidate regions B (C9: 65,000-80,000) and C (C11:615,000-620,000). Here the haplotypes driving adaptation in the Burke et al.^20^ study actually decrease in frequency in most instances.

**Table 5.** Comparison of putatively adaptive haplotypes from Burke et al. (2014) with observed haplotype frequency changes in synthetic populations used in this study. While both studies employed similar protocols Burke et al. (2014) implemented 18 outcrossing cycles while we only implemented 12. The 2014 study is used as a point of reference here because it features 12-fold replication, while in the present study features no within-treatment replication. We specifically targeted the window containing the most significant marker in each region based on the Burke et al. (2014) results. Notable (> 10%) increases in focal haplotype frequency are shown in green.

| Candidate region | Haplotype change in Burke et al. (2014) |  | Haplotype change in populations of the present study |  |  |  |  |  |
| --- | --- | --- | --- | --- | --- | --- | --- | --- |
|  | Driving haplotype | Mean increase | S4 | K4 | S8 | K8 | S12 | K12 |
| A- C07:950000-975000 | Y12 | +0.23 | +0.49 | +0.29 | +0.36 | +0.09 | +0.16 | +0.66 |
| B- C09:65000-80000 | DBVPG6044 | +0.24 | +0.70 | -0.01 | -0.20 | -0.16 | +0.15 | +0.49 |
| C- C11:615000-620000 | DBVPG6044 | +0.29 | -0.16 | +0.01 | -0.06 | -0.15 | -0.11 | +0.49 |
| E- C16:445000-460000 | Y12 | +0.59 | +0.29 | +0.51 | +0.11 | +0.93 | +0.50 | +0.98 |

### Phenotypic characterization of experimental populations

Sporulation efficiencies were estimated in all recombinant populations initially, and after 12 cycles of outcrossing (Supplementary Table S3). We find that population K12 at the end of the experiment (after 12 outcrossing cycles) has the lowest sporulation efficiency at ~10%, while all other populations, including the initial K12 population, had sporulation efficiencies exceeding 30%. Excluding K12 cycle 12, estimates range from 30% to 55% with a mean of 38%. Looking across these estimates, we find no clear relationships of sporulation efficiency to either: i) number founders used; ii) K- versus S-type build strategy; or iii) cycle 0 versus cycle 12. But, variation among assays of biological replicates of the same population was high enough (see standard deviations provided in Supplementary Table S3), our ability to make definitive comparisons between these estimates is limited. As such, we can only say that sporulation efficiency in K12 cycle 12 is greatly reduced when compared to the other recombinant populations. We also assessed growth rates in rich media for all 12 founder strains, and the recombinant populations initially and after 12 cycles of outcrossing. We observe variation in the founder strains with doubling times ranging from ~1 hour to ~1.3 hours, but we see no obvious relationships between founders used to create a specific recombinant population and its initial growth rate (Supplementary Table S4). For instance, two of our slowest growing founder strains, YJM975 and 273614N, are only used when creating K12 and S12. However, the 12-founder populations do not grow more slowly than the 4- or 8- founder populations. We do see evidence of a trend where the recombinant populations grow more slowly at cycle 12 than they did initially, but changes are not particularly large (ranging from −0.01 to −0.22 hours with a mean of −0.10).

## Discussion

### Careful crossing of founder strains increases SNP-level variation

Here we primarily sought to assess how crossing strategy - pairwise crossing of founder strains versus mixing populations in equal proportion - impacted levels of SNP and haplotype variation. We considered both the total number of polymorphic sites and levels of heterozygosity at those sites in synthetic populations created using these two approaches. We consistently found that populations generated by imposing a round of careful crossing have more polymorphic sites (Table 2), and less variation is lost at those sites over time, compared to populations created by simply mixing founders in equal proportion (Fig. 2 and Table 3). We also found that increasing the number of founder strains used to create a given synthetic population also generally resulted in higher levels of SNP variation. However, the total percentage of potential polymorphic sites lost was typically higher when more populations were used (Table 2). And combining the K-type crossing strategy with a large number of founders resulted in dramatic skews in the site frequency spectrum and losses in SNP variation (Fig. 1). Lastly, we find no evidence that specific sets of variants are favored in one crossing strategy versus another when the same founders are used (Supplementary Fig S3). Instead, we suggest that differences in outcomes between crossing strategies are most likely due to strain-specific differences in mating efficiencies and reproductive output (i.e. when populations are simply mixed together, the most compatible strains dominate).

Overall, these findings have led us to two general recommendations. First, if one endeavors to produce populations with the highest possible levels of total SNP variation, many founders and a crossing strategy that involves at least one round of pairwise crossing should be considered. However, if one’s goal is to preserve as much of the variation in founding genotypes as possible, a crossing strategy with fewer founders might be more desirable.

### Careful crossing of founder strains results in more balanced haplotype representation

Haplotype frequency estimates for our synthetic populations suggest that a strategy involving pairwise crosses results in better representation and maintenance of founder genotypes (Fig. 4, Supplementary Figs. S5-S18, and Table 4). It is clear than when haploid strains are simply pooled, there is potential for subsets of the founder strains to dominate and skew haplotype representation in the resulting populations. As such, we recommend a careful (i.e. pairwise) crossing strategy when seeking to maximize founder haplotype representation. In addition, using fewer founder haplotypes also generally results in more even representation among them, though we would not necessarily recommend against using more founders unless it is crucial to achieve an even haplotype representation. In fact, we would argue that there are empirical benefits to a more varied distribution of haplotype frequencies segregating within a synthetic population. Specifically, in an E&R context, such a population creates opportunities to observe selection on both rare and common haplotypes, and the relative contributions each to the evolution of quantitative traits remains an unresolved question (e.g. Bloom et al.^41^).

What might be driving the extreme pattern of haplotype loss we observe in the K-type populations compared to their S-type counterparts? Here we outline two possibilities, using the K4 and S4 populations to illustrate. In the former, we find that the YPS128 and DBVPG6044 haplotypes are almost entirely missing while in the latter all founder haplotypes are evenly represented. We think that the most obvious mechanism underlying this pattern is the reproductive capacity of the founding strains; in other words, if particular strains inefficiently sporulate and/or mate, or are not compatible with other strains, haplotype loss should result. The two missing haplotypes in K4 appear to have been lost immediately after the population’s creation (Supplementary Figure S11), which would support the idea that these two genotypes are simply disadvantageous under environmental circumstances which require cells to sexually reproduce to survive. A second, non-mutually exclusive possibility that could drive the haplotype loss we observed is the emergence of an asexual diploid lineage that can evade our forced outcrossing protocols and become the majority genotype through clonal interference. Others working with recombinant *S. cerevisiae* have reported evidence of such “cheater” lineages (Linder et al.^42^; note: their crossing protocol, described in Linder et al.^23^, represents a middle ground between our K and S strategy). Since we observe intermediate levels of haplotype representation in the K4 population mid-way through the experiment (Supplementary Figure S12), this suggests that if a diploid cheater lineage emerged, this happened over a long evolutionary timescale; in other words, this cannot explain the early haplotype losses we observed in the population, but it might explain long-term loss of haplotype diversity. At the same time, one would expect a expect a cheater genotype to take over very rapidly in the population, which we did not observe. In summary, we think our experiments provide compelling evidence that differences in sexual reproduction between strains will lead to haplotype loss in the K-type populations, and that this loss can be prevented by using an S-type strategy. While the evolution of cheating could reasonably occur in any of our populations, and also result in haplotype loss, we report no strong evidence that this is more likely to happen in K-type versus S-type populations.

### Evidence for adaptation to outcrossing

While we interpret the differences between the different crossing strategies (i.e. between all S-type and K-type populations) as being primarily driven by initial differences in sporulating and mating efficiencies between strains and subsequent genetic drift, selection may also play a role. For instance, we do find evidence that two specific regions of the genome exhibit similar dynamics in haplotype frequencies, compared to similar prior work; Burke et al.^20^ previously implicated these regions as potentially driving adaptation for frequent outcrossing. We also compared our results to those of Linder et al.^23^ which features synthetic populations constructed using the same founder strains present in this study. However, here we did not find it was the case that haplotypes most common in their populations on average are also the most common in our populations. For instance, Y12 is a common haplotype across all of our populations (Table 4) with mean genome-wide frequencies ranging from 0.2 to ~0.5, but it does not exceed a frequency of 0.06 in either of their synthetic populations (*cf*. Table 3 of Linder et al.^20^). Similarly, YPS128 which has a mean frequency of 0.41 in one of the their populations appears at frequencies below 0.11 in all of our populations except S4. That being said, it is worth noting that while there is overlap in the strains used between these studies, maintenance protocols are different, and their populations include several strains absent in our study. So, we can only say that no common founders appear to universally favored when selection for frequent outcrossing is imposed.

Our analysis comparing initial SNP frequencies to those observed after 12 rounds of outcrossing in each population also yields possible evidence for adaptation, though our interpretation here is cautious. We do observe regions that produce peaks in significance in some populations that overlap regions described by other studies to underlie traits related to outcrossing (e.g. Fig. 3D; peak in C16 in K8). However, peaks are rarely recapitulated across two or more populations (Fig. 3) and in some populations we observe no clearly defined peaks at all (e.g. Fig. 3B; population K4). As such, our results are at best suggestive. But, we find it notable that a crossing strategy using 8 founding haplotypes leads to the most overlap with other candidate regions identified in the literature. While this experiment has limited ability to associate particular genomic regions and phenotypes related to outcrossing, this observation leads us to speculate that using an intermediate number of founding haplotypes (e.g. 8) may lead to an increased ability to localize candidate regions in an E&R experiment.

Given that any adaptation we did observe can only be ascribed to the outcrossing process and associated handling protocols, we conclude that there is likely a great deal of value in allowing newly-established synthetic populations to experience several cycles of outcrossing – this can also be thought of as laboratory domestication – before any sort of new selective pressure is imposed. To be explicit, if some other selection pressure was intentionally imposed on our populations immediately following cycle 0, it would be very difficult to dissect the specific genetic changes that might occur due to that pressure, other laboratory handling steps, or general selection for reproduction via outcrossing.

### Crossing strategy and number of founder strains does not obviously impact sporulation or growth rate

We assayed sporulation efficiencies and growth rates in recombinant populations as these are both important fitness related characters that may be impacted by crossing strategy, or that might respond to the selection imposed by many cycles of forced outcrossing. Looking at sporulation efficiencies (Supplementary Table S3), we do not find any obvious associations between these estimates and a particular crossing strategy. We also find no clear evidence that sporulation efficiency increases over the course of the experiment, which is somewhat surprising given the forced outcrossing that defines our maintenance protocol. The only major finding that emerges is that sporulation efficiency is much lower in K12 cycle 12 than what we observe in any other population, including the initial K12 population. We speculate that this is perhaps related to the wholesale loss of genetic variation in K12, or this may indicate that asexual diploid lineages representing only a fraction of total variation among founders have risen to prominence. However, our ability to make any definitive statements about how crossing strategy or number of founders shapes sporulation efficiency as a life-history trait is limited.

Comparing growth rates of founder strains to recombinant populations (Supplementary Table S4) similarly does not reveal clear evidence linking growth rates in founder strains to either K or S-type populations (i.e. recombinant populations have similar doubling times regardless of which strains were used or how they were combined). It is also not the case that our fastest growing founders are better represented when looking haplotype estimates or vice versa. There is a consistent trend of slower growth rates in cycle 12 versus cycle 0 for the recombinant populations. Differences are small, but this is still perhaps suggestive of some sort of trade-off between growth and other fitness characters as populations adapt to our maintenance protocols. In other words, it is conceivable that the demands for high levels of outcrossing might result in populations that invest more in sexual reproduction, and less in budding. However, we find no consistent patterns when comparing difference between S and K strategies or number of founders. As such, we find no evidence that these factors are shaping growth rates in the recombinant populations.

### Conclusions

The results of simulated E&R studies in which populations are sexually reproducing and adaptation is driven by standing genetic variation have led to general experimental design recommendations that maximize genetic variation in the ancestral population, specifically by increasing the number of starting haplotypes^7–8^. Here, we provide empirical results that also these recommendations. Across the metrics we examined, we consistently find that a crossing strategy involving careful pairwise crosses leads to populations with more standing genetic variation than those produced by simply mixing founder genotypes in equal proportion. As such, using this sort of strategy would be our primary recommendation for researchers aiming to establish recombinant populations from clonal strains or isogenic lines for use in E&R studies. Using more founders also results in greater total levels of genetic variation but comes at the cost of maximizing representation of all possible alleles from a given set of founders. So, the number of founders to used should be chosen based upon the specific goals of an E&R study, and the questions it hopes to test. Finally, our results should be placed in the context of yeast biology. Meaning, given the high degree of genetic differentiation between our yeast strains and the variation among them for mating and sporulation efficiencies, the patterns we observe are likely more extreme than what might be expected when creating a *Drosophila* synthetic population using either the K-type or S-type approach. However, it is certainly conceivable that mating preferences and genetic incompatibilities among isogenic *Drosophila* lines could impact levels of genetic variation and haplotype representation when they are crossed. Our findings therefore perhaps warrant consideration even when creating synthetic populations in non-yeast systems.

## Data Availability

The raw sequence files generated over the course of this project are available through NCBI SRA (BioProject ID: PRJNA732717). Core data files (tables with SNP and haplotype frequencies, results of statistical analysis, etc.) are available through Dryad (doi:10.5061/dryad.g79cnp5qg). Script used to process raw data and perform SNP calling are available through Github (https://github.com/mollyburke/Burke-Lab-SNP-calling-pipeline), as are the core scripts necessary to reproduce our results (https://github.com/mphillips67/Build-Paper).

## Acknowledgements

We thank undergraduate researcher Julia Reinsch for her help with sporulation assays, and Dr. Anthony D. Long for his feedback on the haplotype estimation analysis. We also thank Oregon State University’s Center for Genome Research and Biocomputing for use of their computational and sequencing resources. This work was supported by startup funds provided to M.K.B. by College of Science at Oregon State University, and M.A.P. was supported by a National Science Foundation Postdoctoral Fellowship (NSF 1906246).

## Author Contributions

M.K.B. and I.C.K. conceived of the project. I.C.K. performed the lab work necessary to generate the genomic data sets featured in this study, and K.M.M. and S.K.T. generated all phenotypic data and results. M.A.P. and M.K.B. formulated the analytic strategy for the genomic data, and M.A.P. performed all major genomic analyses. M.A.P. and M.K.B. wrote the manuscript.

## Notes

### Competing Interest Statement

The authors have declared no competing interest.

## References

1. Manolio, T.A. et al. Finding the missing heritability of complex diseases. Nature 461, 747–753 (2009).

2. Boyle, E.A., Li, Y.I. & Pritchard, J.K. An expanded view of complex traits: from polygenic to omnigenic. Cell 169:1177–1186 (2017).

3. Schlötterer, C., Tolber, R., Kofler, R., & Nolte, V. Sequencing pools of inidviduals – mining genome-wide polymorphism data without big funding. Nat. Rev. Genet. 15, 749–763 (2014).

4. Long A.D., Liti, G., Luptak, A. & Tenaillon O. Elucidating the molecular architecture of adaptation via evolve and resequence experiments. Nat. Rev. Genet., 16, 567–582 (2015).

5. Bailey, S.F. & Bataillon, T. Can the experimental evolution programme help us elucidate the genetic basis of adaptation in nature? Mol. Ecol. 25, 203–216 (2016).

6. Burke, M.K. How does adaptation sweep through the genome? Insights from long-term selection experiments. Proc. Roy. Soc. B., 279, 5029–5038 (2012).

7. Baldwin-Brown, J.G., Long, A.D. & Thornton, K.R. The power to detect quantitative trait loci using resequenced, experimentally evolved populations of diploid, sexual organisms. Mol. Biol. Evol. 31, 1040–1055 (2014).

8. Kofler, R. & Schlötterer, C. A guide for the design of evolve and resquencing studies. Mol. Biol. Evol. 31, 473–482 (2014).

9. Vlachos, C. & Kofler, R. Optimizing the power to identify the genetic basis of complex traits with evolve and resequence studies. Mol. Biol. Evol. 35, 2890–2905 (2019).

10. Burke, M.K. et al. Genome-wide analysis of a long-term evolution experiment with *Drosophila*. Nature 467, 587–590 (2010).

11. Turner, T.L., Stewart, A., Fields, A.T., Rice, W.R. & Tarone, A.M. Population-based resequencing of experimentally evolved populations reveals the genetic basis of body size variation in *Drosophila melanogaster*. PLoS Genet., 7, e10001336 (2011).

12. Orozco-ter Wengel, P. et al. Adaptation of *Drosophila* to a novel laboratory environment reveals temporally heterogeneous trajectories of selected traits. Mol. Ecol. 21, 4931–4941 (2012).

13. Tobler, R. et al. Massive habitat-specific genomic response in D. *melanogaster* populations during experimental evolution in hot and cold environments. Mol. Biol. Evol. 31, 364–375 (2014).

14. Huang, Y., Wright, S.I. & Agrawal, A.F. Genome-wide patterns of genetic variation within and among alternative selective regimes. PLoS Genet. 10, e1004527 (2014).

15. Jha, A.R. et al. Whole-genome resequencing of experimental populations reveals polygenic basis of egg-size variation in *Drosophila melanogaster*. Mol. Biol. Evol. 32, 2616–2632 (2015).

16. Franssen, S.U., Nolte, V., Tobler, R. & Schlötterer C. Patterns of linkage disequilibrium and long range hitchhiking in evolving experimental *Drosophila* melanogaster populations. Mol. Biol. Evol. 32, 495–509 (2015).

17. Graves, J.L. et al. Genomics of parallel experimental evolution in Drosophila. Mol. Biol. Evol. 34, 831–842 (2017).

18. Phillips, M.A. et al. Effects of evolutionary history on genome wide and phenotypic convergence in Drosophila populations. BMC Genomics 19, 743 (2018).

19. Barghi, N. et al. Genetic redundancy fuels polygenic adaptation in *Drosophila*. PLOS Biol. 17, e3000128 (2019).

20. Burke, M.K., Liti, G. & Long, A.D. Long standing genetic variation drives repeatable experimental evolution in outcrossing populations of *Saccharomyces cerevisiae*. Mol. Biol. Evol. 32, 3228–3239 (2014).

21. Phillips, M.A., Kutch, I.C., Long, A.D. & Burke, M.K. time sampling in an eolve-and- resequence experiment with outcrossing *Saccharomyces cerevisiae* reveals multiple paths of adaptive change. Mol. Ecol., 29, 4898–4912 (2020).

22. Cubillos, F.A et al. High-resolution mapping of complex traits with a four-parent advanced intercross yeast population. Genetics 195, 1141–1155 (2013).

23. Linder, R.A., Majumbder, A., Chakraborty, M. & Long, A.D. Two synthetic 18-wayoutcrossed populations of diploid budding yeast with utility for complex trait dissection. Genetics 215, 323–342 (2020).

24. King, E.G. et al. (2012). Genetic dissection of a model complex trait using the Drosophila synthetic population resource. Genome Res., 22, 1558–1566.

25. Teotónio, H., Carvalho, S., Manoel, D., Roque, M. & Chelo, I.M. Evolution of outcrossing in experimental populations of *Caenorhabditis elegans*. PLoS One, 7, e35811 (2012)

26. Michalak, P., Kang, L., Schou, M.F., Garner, H. & Loeschke, V. Genomic signatures of experimental adaptive radiation in Drosophila. Mol. Ecol. 28, 600–614 (2018).

27. Barghi. N. & Schlötterer, C. Shifting the paradigm in evolve and resequence studies, from analysis of single nucleotide polymorphism to selected haplotype blocks. Mol. Ecol. 28, 521–524 (2019).

28. Nouhaud, P., Tobler, R. Nolte, V. & Schlötterer, C. Ancestral population reconstitution from isofemale lines as a tool for experimental evolution. Eco. Evol. 6, 7169–7175 (2016).

29. Harbison, S.T., Negron Serrano, Y.L., Hansen, N.F. & Lobell, A.S. Selection for long and short sleep duration in *Drosophila melanogaster* reveals the complex genetic network underlying natural variation in sleep. PLoS Genet. 13, e1007098 (2017).

30. Cubillos, F.A., Louis, E.J. & Liti, G. Generation of a large set of genetically tractable haploid and diploid *Saccharomyces* strains. FEMS Yeast Res. 9, 1217–1225 (2009).

31. Burke, M.K., McHugh, K.M. & Kutch, I.C. Heat shock improves random spore isolation in diverse strains of *Saccharomyces cerevisiae*. Front. Genet. 11: 597482 (2020).

32. Baym, M. et al. Inexpensive multiplexed library preparation for megabase-sizes genomes. PLoS One 10, e0131262 (2015).

33. Mkenna et al. The genome analysis toolkit: a mapreduce framework for analyzing next-generation DNA sequencing data. Genome Res. 20, 1297–1303 (2010)

34. Poplin et al. A universal SNP and small-indel variant caller using deep neural networks. Nat. Biotechnol. 36, 983–987 (2018).

35. Bergström, A. et al. A high-definition view of functional genetic variation from natural yeast genomes. Mol. Biol. Evol. 31, 872–888 (2014).

36. Conway, J.R., Lex, A. & Gehlenborg, N. UpSetR: an R package for the visualization of intersecting sets and their properties. Bioinformatics, 33, 2938–2940 (2017).

37. Beissinger, T.M., Rosa, G.J.M, Kaeppler, S.M., Gianola, D. & de Leon, N. Defining window-boundaries for genomic analyses using smoothing spline techniques. Genet Sel. Evol. 47, 30 (2015).

38. Taus, T., Futschik, A. & Schlötterer, C. Quantifying selection with pool-seq time series data. Mol. Biol. Evol., 34, 3023–3034 (2017).

39. Nei, M. & Tajima, F. DNA polymorphism detectable by restriction endonucleases. Genetics 97, 145–163 (1981).

40. Sprouffske, K. & Wagner, A. Growthcurver: an R package for obtaining interpretable meterics from microbial growth curves. BMC Bioinform. 17, 172 (2016).

41. Bloom, J.S. et al. Rare variants contribute disproportionately to quantitative trait variation in yeast. eLife 8, e49212 (2019).

42. Linder, R.A. et al. Adaptation in outbred sexual yeast is repeatable, polygenic, and favors rare haplotypes. bioXiv, doi.org/10.1101/2021.08.27.457900 (2021).

