## Supplemental Figures and Tables for "Crossing design shapes patterns of genetic variation in synthetic recombinant populations of *Saccharomyces cerevisiae*"

**Table of Contents**

| | | page |
|---|---|---|
| **Figure S1** | Strains used in this study. | **3** |
| **Figure S2** | Crossing design used to create S4 (A), S8 (B), and S12 (C). | **4** |
| **Figure S3** | UpSet plots illustrating similarities in genetic variation in recombinant populations. | **5** |
| **Figure S4** | Expected site frequency spectra for initial populations created using four (A), eight (B) and twelve founder strains (C). | **6** |
| **Figure S5** | Haplotype frequency estimates for population S4 after 12 cycles of outcrossing. | **7** |
| **Figure S6** | Haplotype frequency estimates for population K4 after 12 cycles of outcrossing. | **8** |
| **Figure S7** | Haplotype frequency estimates for population S8 after 12 cycles of outcrossing. | **9** |
| **Figure S8** | Haplotype frequency estimates for population K8 after 12 cycles of outcrossing. | **10** |
| **Figure S9** | Haplotype frequency estimates for population S12 after 12 cycles of outcrossing. | **11** |
| **Figure S10** | Haplotype frequency estimates for population K12 after 12 cycles of outcrossing. | **12** |
| **Figure S11** | Initial haplotype frequency estimates for population K4. | **13** |
| **Figure S12** | Haplotype frequency estimates for population K4 after 6 cycles of outcrossing. | **14** |
| **Figure S13** | Initial haplotype frequency estimates for population S4. | **15** |
| **Figure S14** | Haplotype frequency estimates for population S4 after 6 cycles of outcrossing. | **16** |
| **Figure S15** | Initial haplotype frequency estimates for population K8. | **17** |
| **Figure S16** | Haplotype frequency estimates for population K8 after 6 cycles of outcrossing. | **18** |
| **Figure S17** | Initial haplotype frequency estimates for population S8. | **19** |
| **Figure S18** | Haplotype frequency estimates for population S8 after 6 cycles of outcrossing. | **20** |
| **Table S1** | Average genome-wide coverage at SNPs identified in each synthetic recombinant population. | **21** |
| **Table S2** | Mean genome-wide haplotype diversity for each synthetic recombinant population after 12 cycles of outcrossing. | **22** |
| **Table S3** | Sporulation efficiencies in recombinant populations. | **23** |
| **Table S4** | Comparative growth rate estimates in recombinant populations and parental strains. | **24** |

### Supplementary Figures

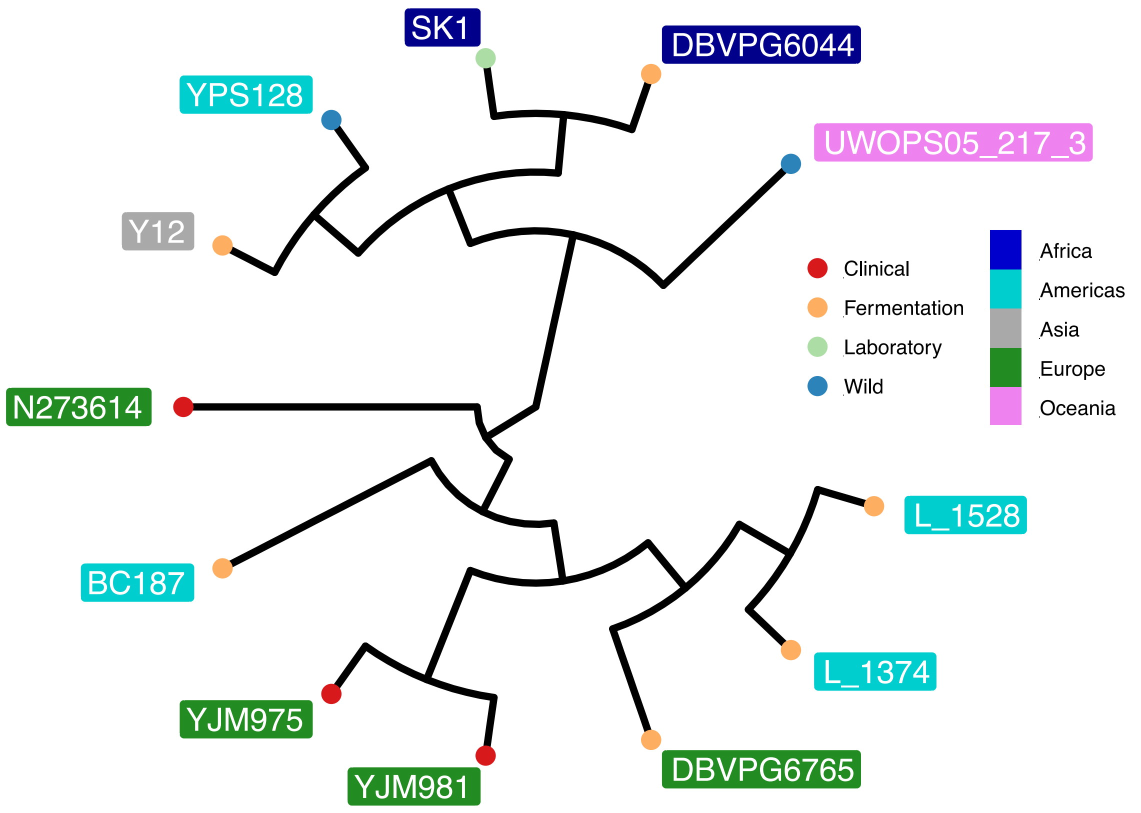

**Supplementary Figure S1**. Strains used in this study. The unrooted phylogeny illustrates the general evolutionary relationships between the strains used in this work, as well as their geographical (colored boxes) and contextual (colored circles) origins. This tree was constructed using 180,276 bi-allelic SNPs that we observed to vary across the 12 haploid founder strains sequenced in this study. These SNPs were used to construct a Newick file which was plotted with the R package *ggtree*.

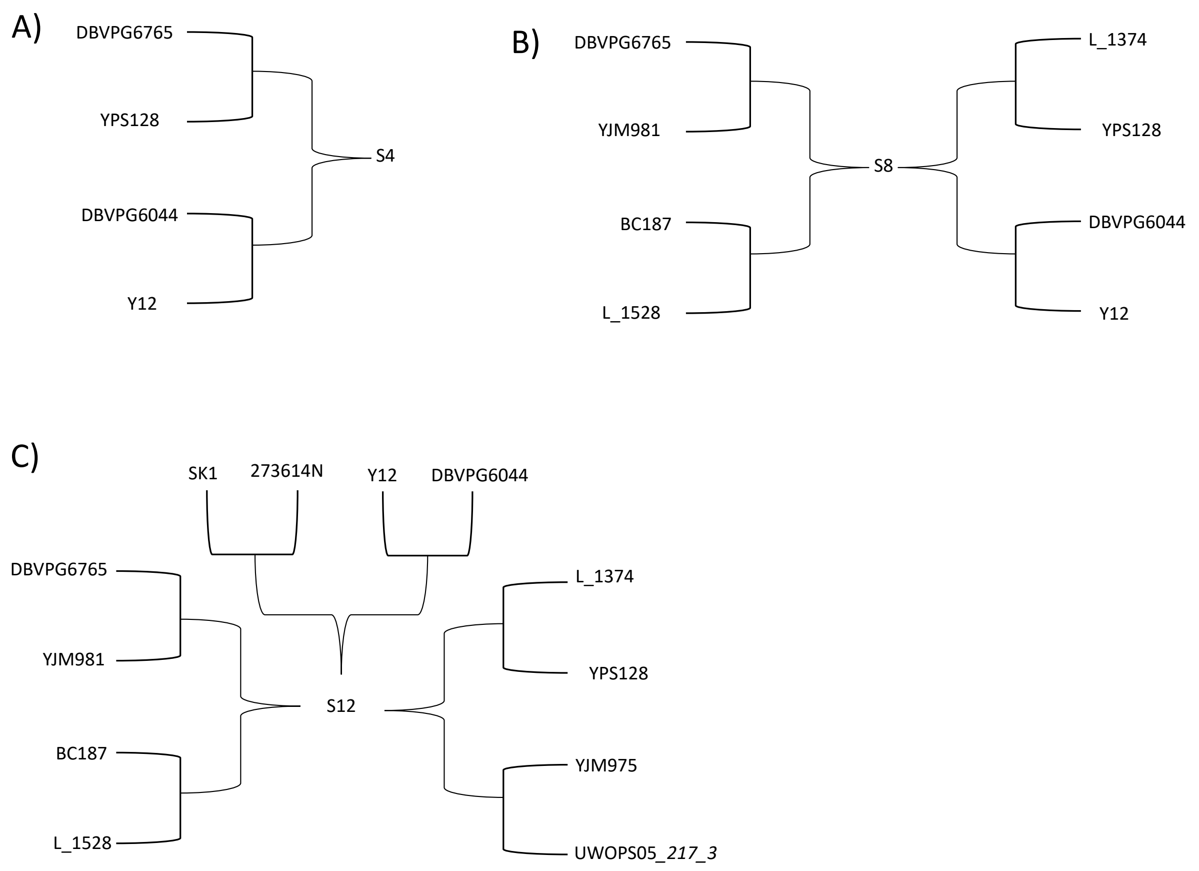

**Supplementary Figure S2.** Crossing design used to create S4 (A), S8 (B), and S12 (C).

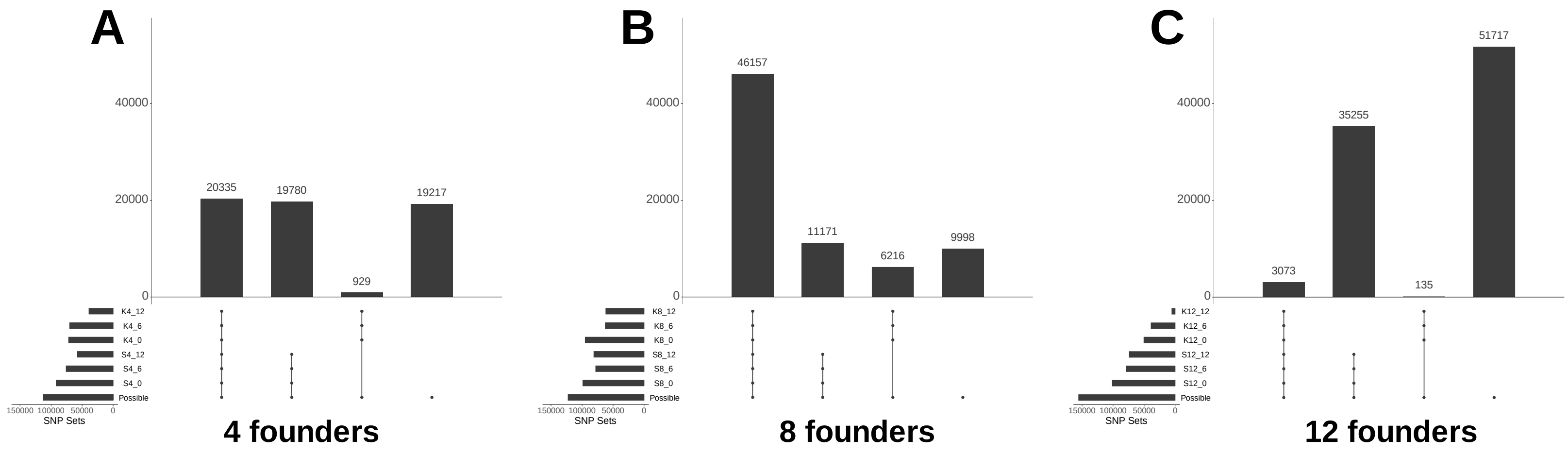

**Supplementary Figure S3.** UpSet plots illustrating similarities in genetic variation in recombinant populations. Plots are organized to compare each pair of S/K populations made using 4 (A), 8 (B), and 12 (C) founders over time. The horizontal bars indicate the size of the SNP sets for each population at each timepoint (t=0, 6, or 12 cycles of outcrossing), and the total possible number of SNPs given the founder strains used to make them. The vertical bars represent distinct intersections between the different highlighted sets (as indicated by the black circles). For a given plot, the first vertical bar represents the number of SNPs found in both K and S-type populations across all timepoints, the second bar the SNPs found in the S-type population across all timepoints but not in the K-type, the third bar the SNPs found across all timepoints in the K-type population but not in the S-type, and the fourth bar the number of possible SNPs that are not present in either the K or S type population at any timepoint (Note: "possible SNPs" refers to sites that have the potential to be polymorphic given the founder used to create a given pair of populations).

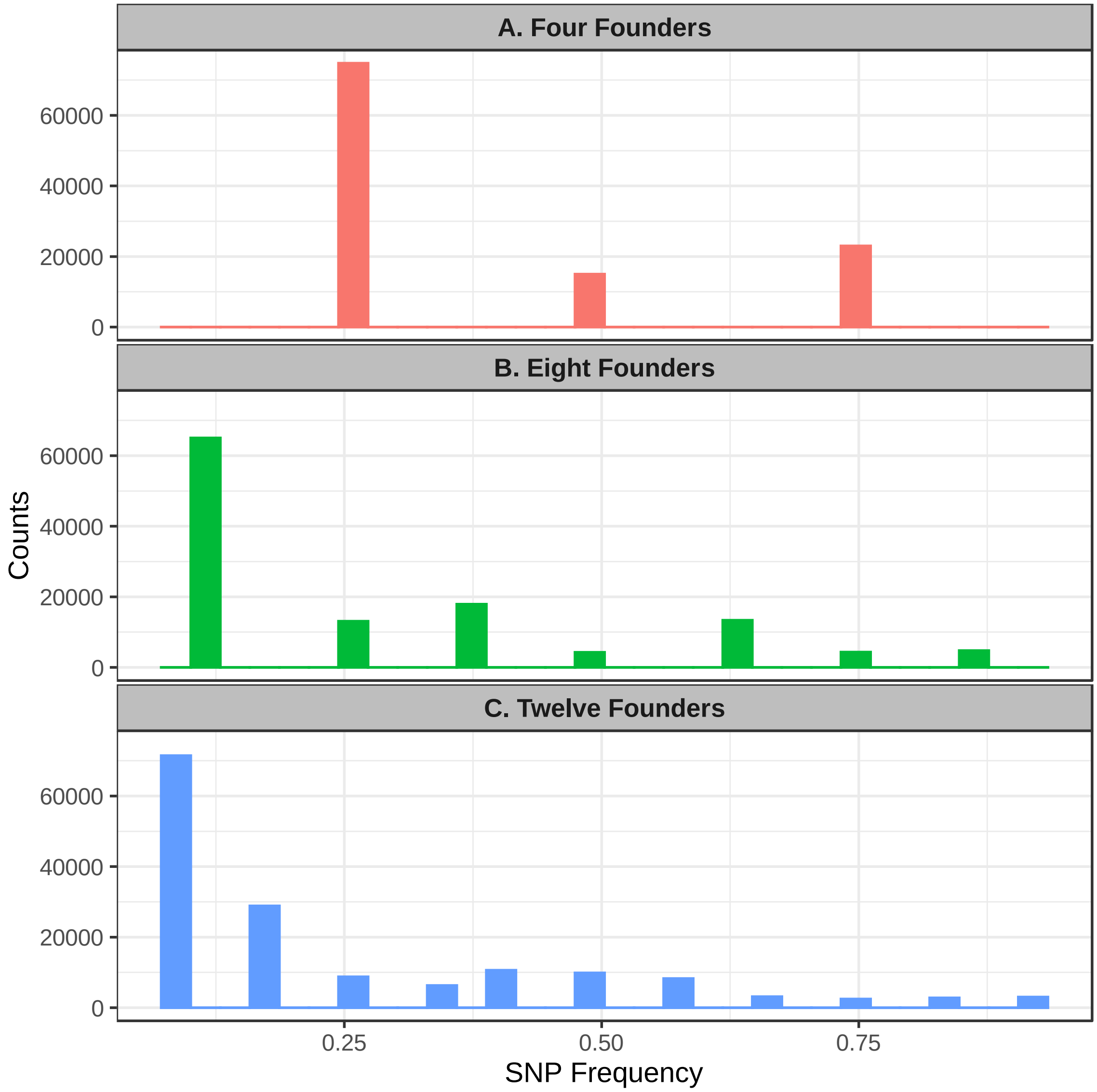

**Supplementary Figure S4**. Expected site frequency spectra for initial populations created using four (A), eight (B) and twelve founder strains (C). For specific founders used in each scenario, see Table 1.

.

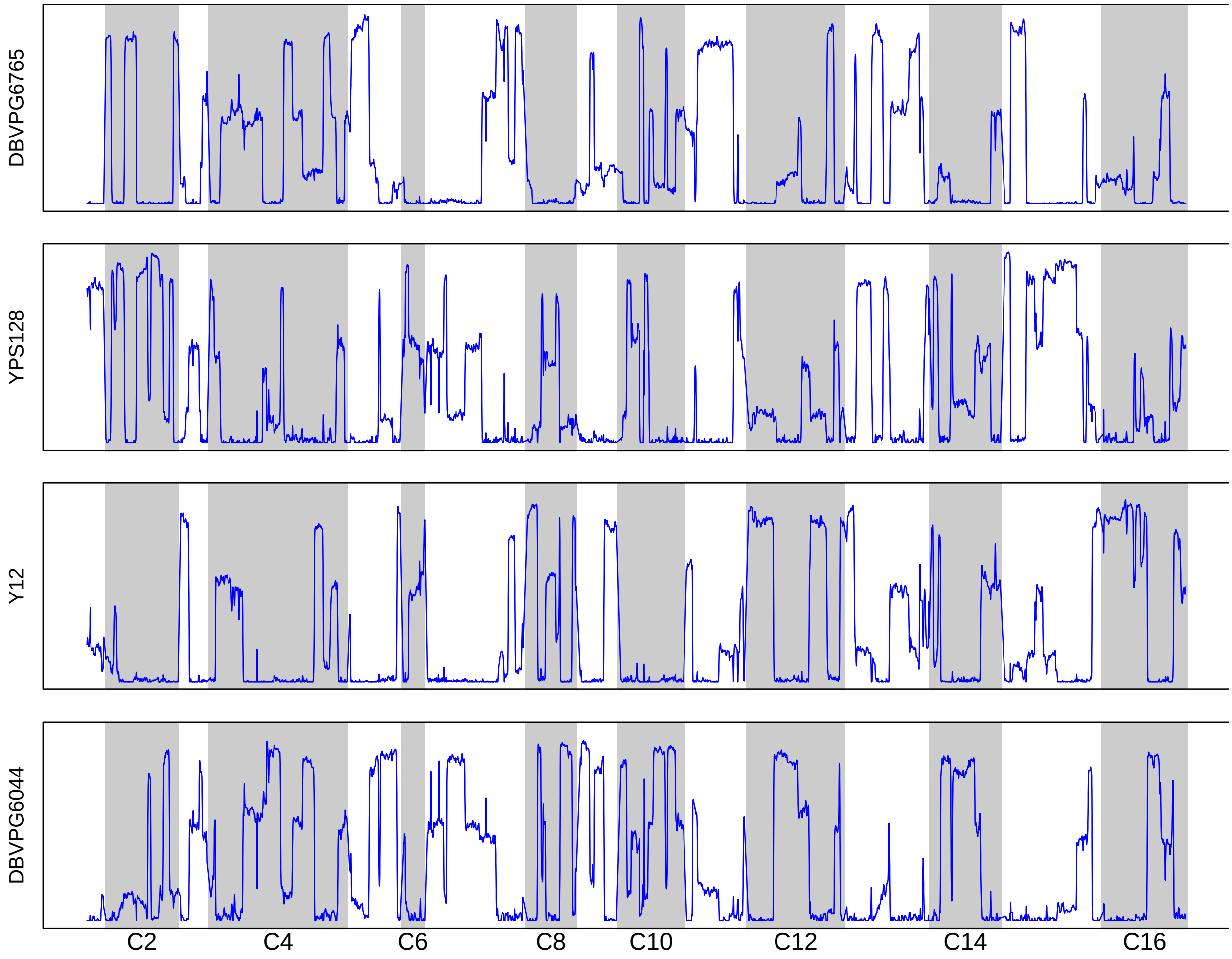

**Supplementary Figure S5.** Haplotype frequency estimates for population S4 after 12 cycles of outcrossing. Frequencies were estimated using sliding windows with a length of 30KB and a 1KB step size. Panels are ordered to match pairings in the first round of crossing (YPS128 was crossed with DBVPG6765, Y12 was crossed with DBVPG6044)

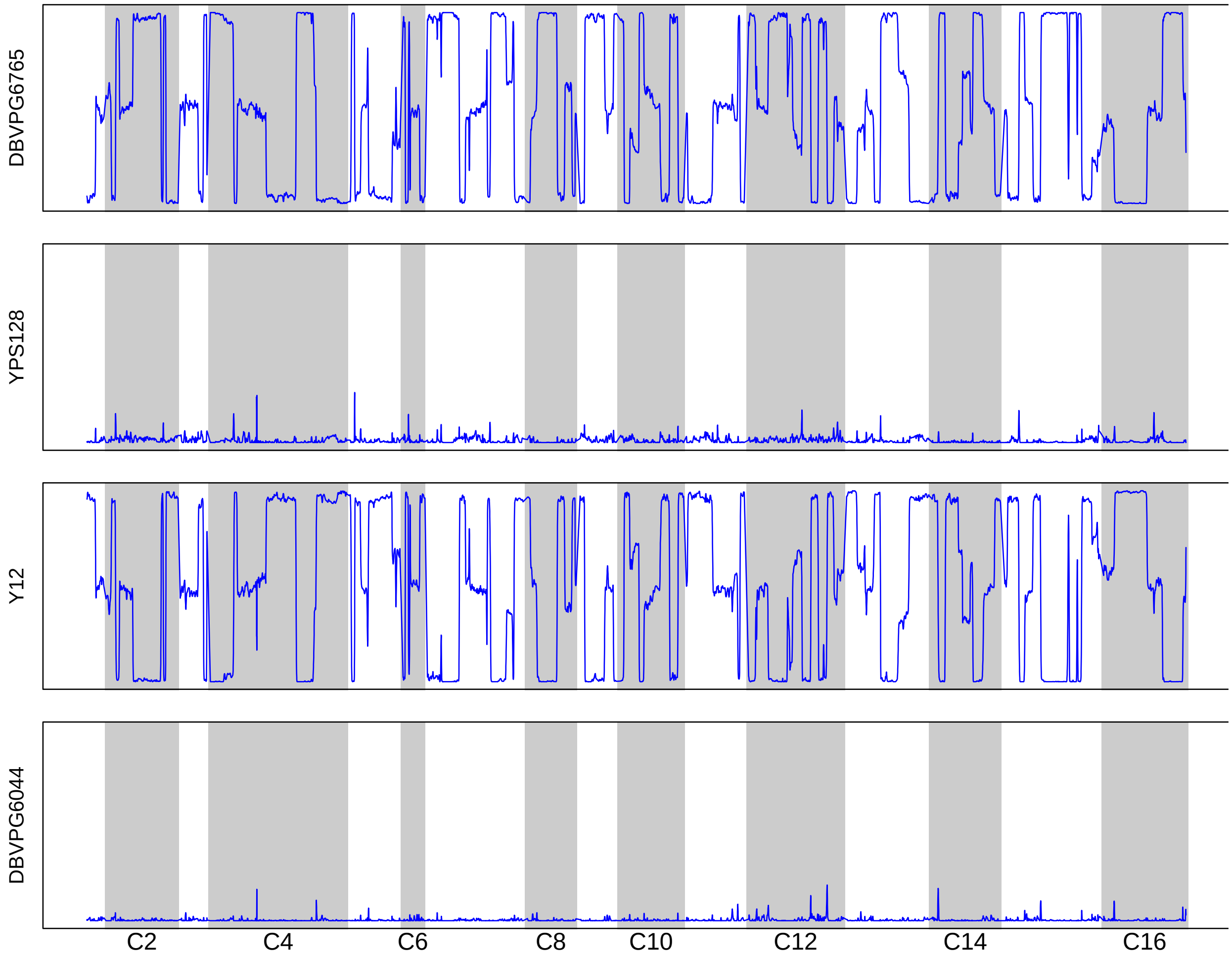

**Supplementary Figure S6.** Haplotype frequency estimates for population K4 after 12 cycles of outcrossing. Frequencies were estimated using sliding windows with a length of 30KB and a 1KB step size.

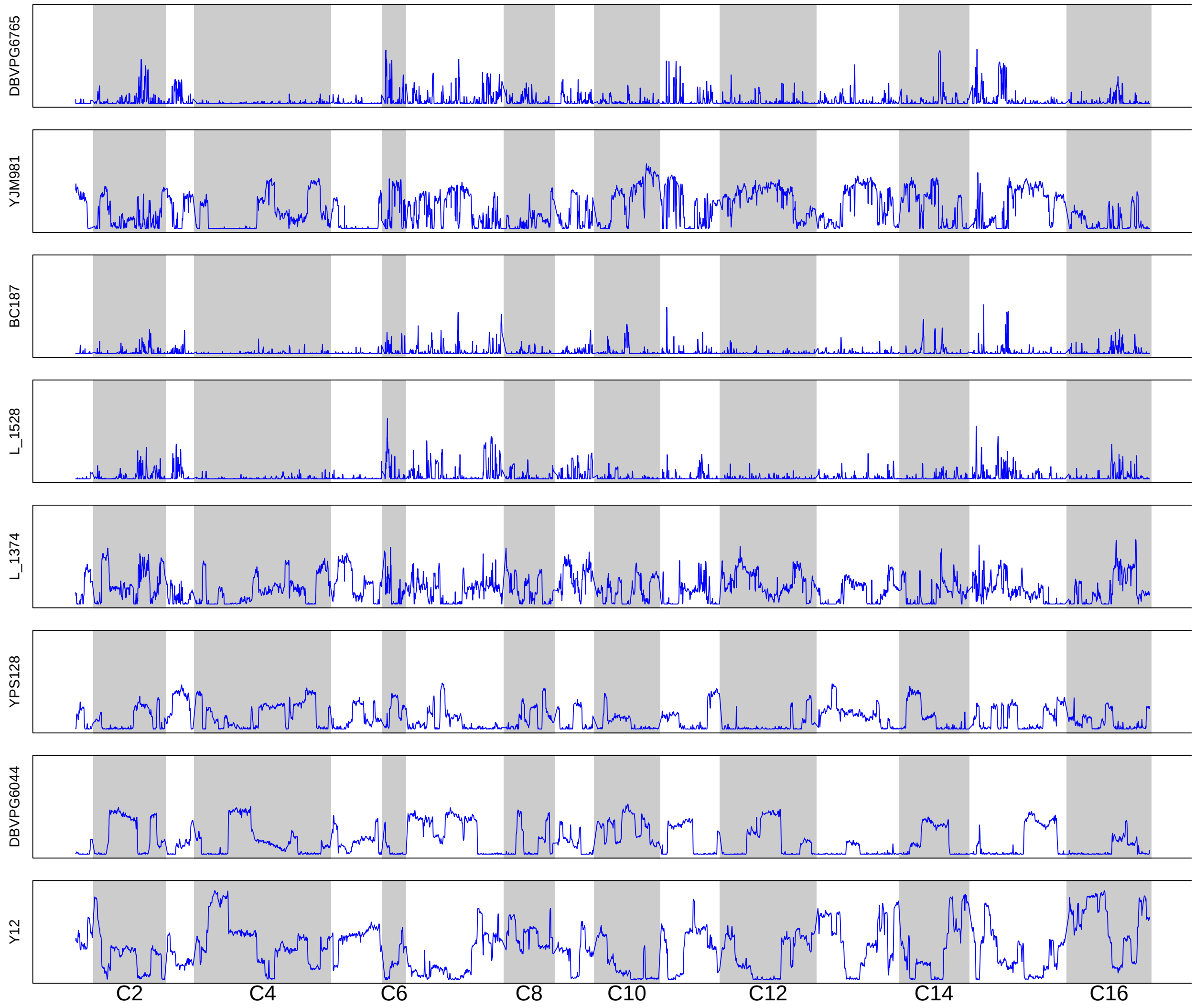

**Supplementary Figure S7.** Haplotype frequency estimates for population S8 after 12 cycles of outcrossing. Haplotype frequencies were estimated using sliding windows with a length of 30KB and a 1KB step size. Panels are ordered to match pairings in the first round of crossing (e.g. YJM981 was crossed with DBVPG6765, etc.).

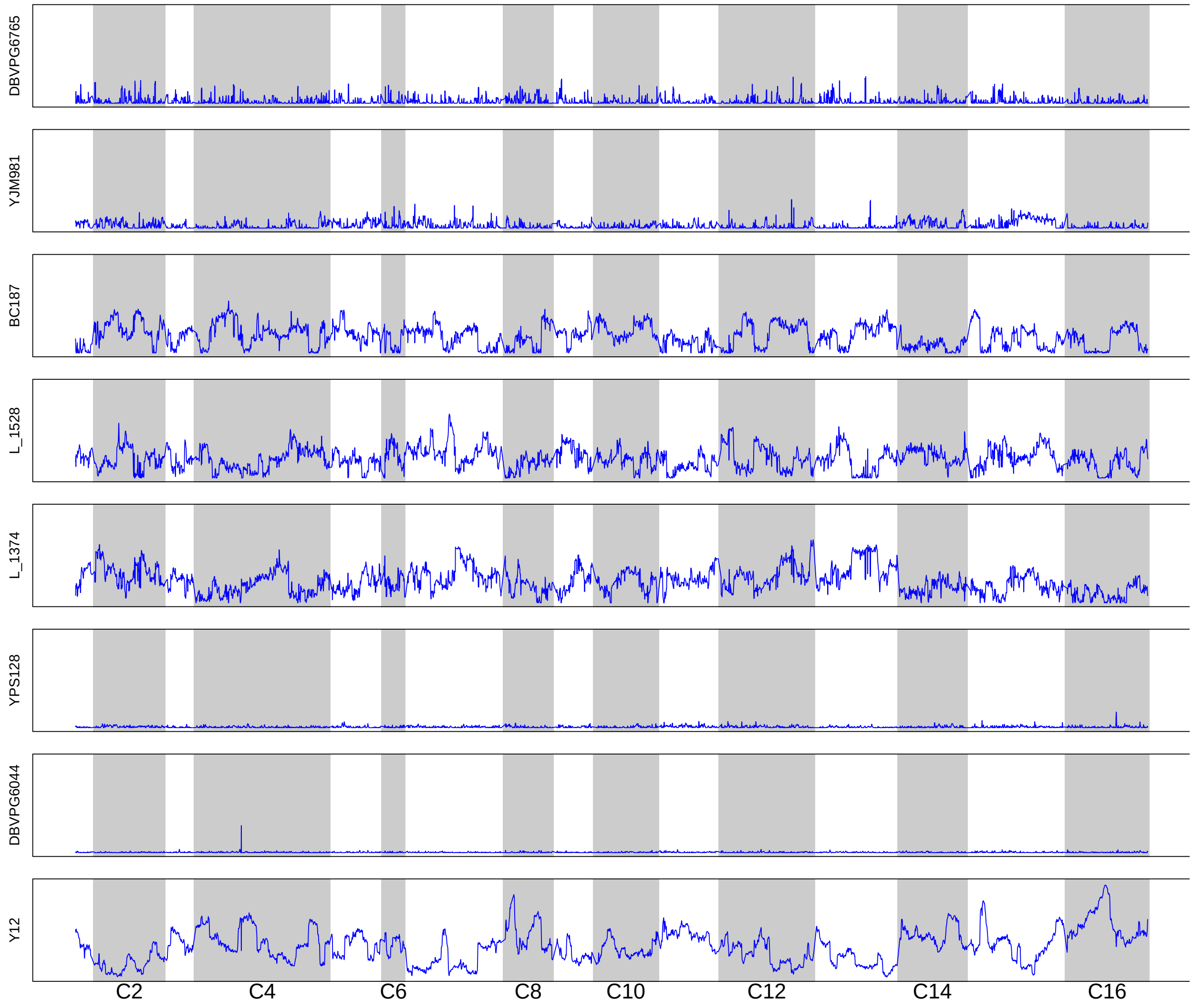

**Supplementary Figure S8.** Haplotype frequency estimates for population K8 after 12 cycles of outcrossing. Haplotype frequencies were estimated using sliding windows with a length of 30KB and a 1KB step size.

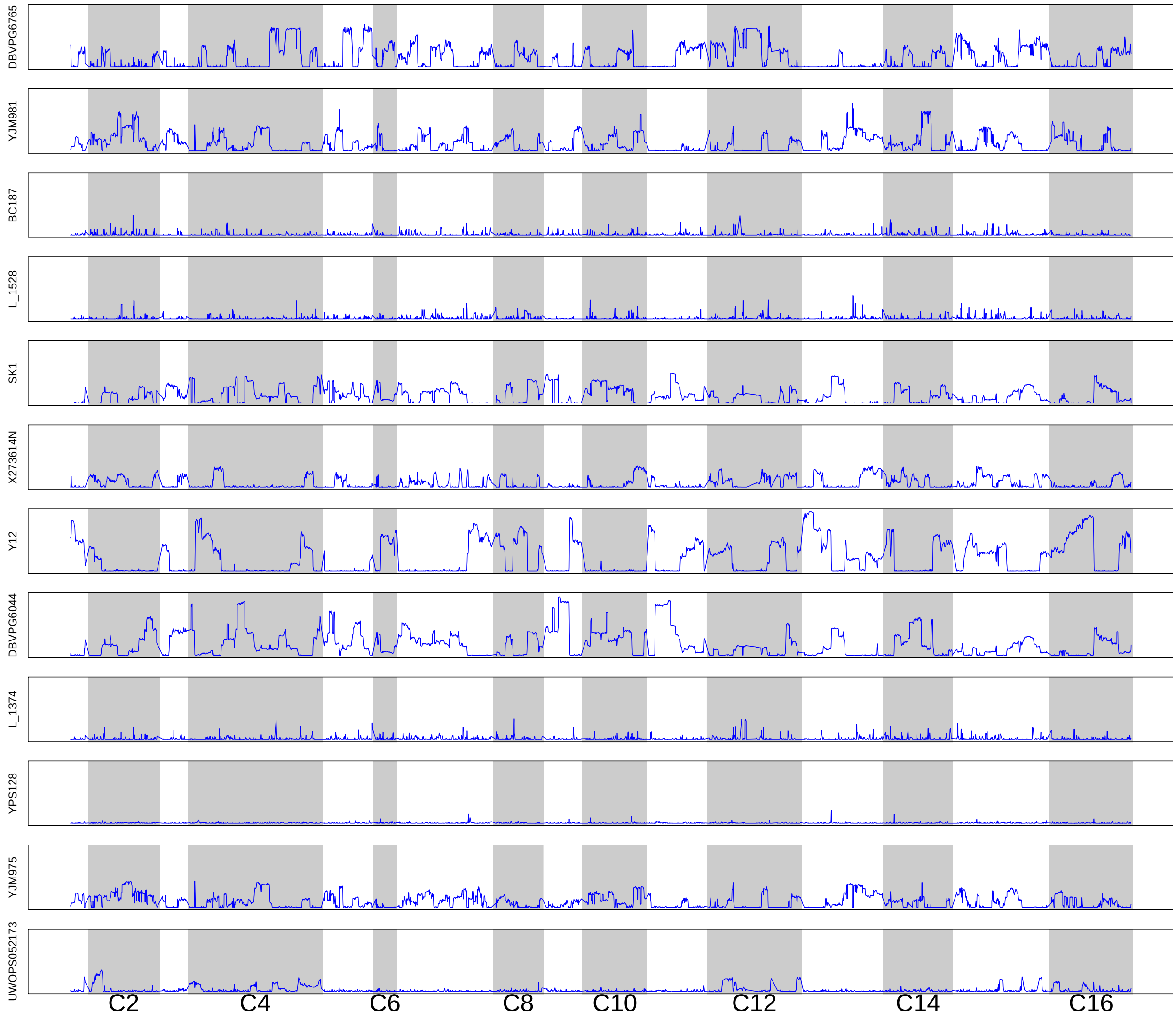

**Supplementary Figure S9.** Haplotype frequency estimates for population S12 after 12 cycles of outcrossing. Haplotype frequencies were estimated using sliding windows with a length of 30KB and a 1KB step size. Panels are ordered to match pairings in the first round of crossing (e.g. YJM981 was crossed with DBVPG6765, etc.).

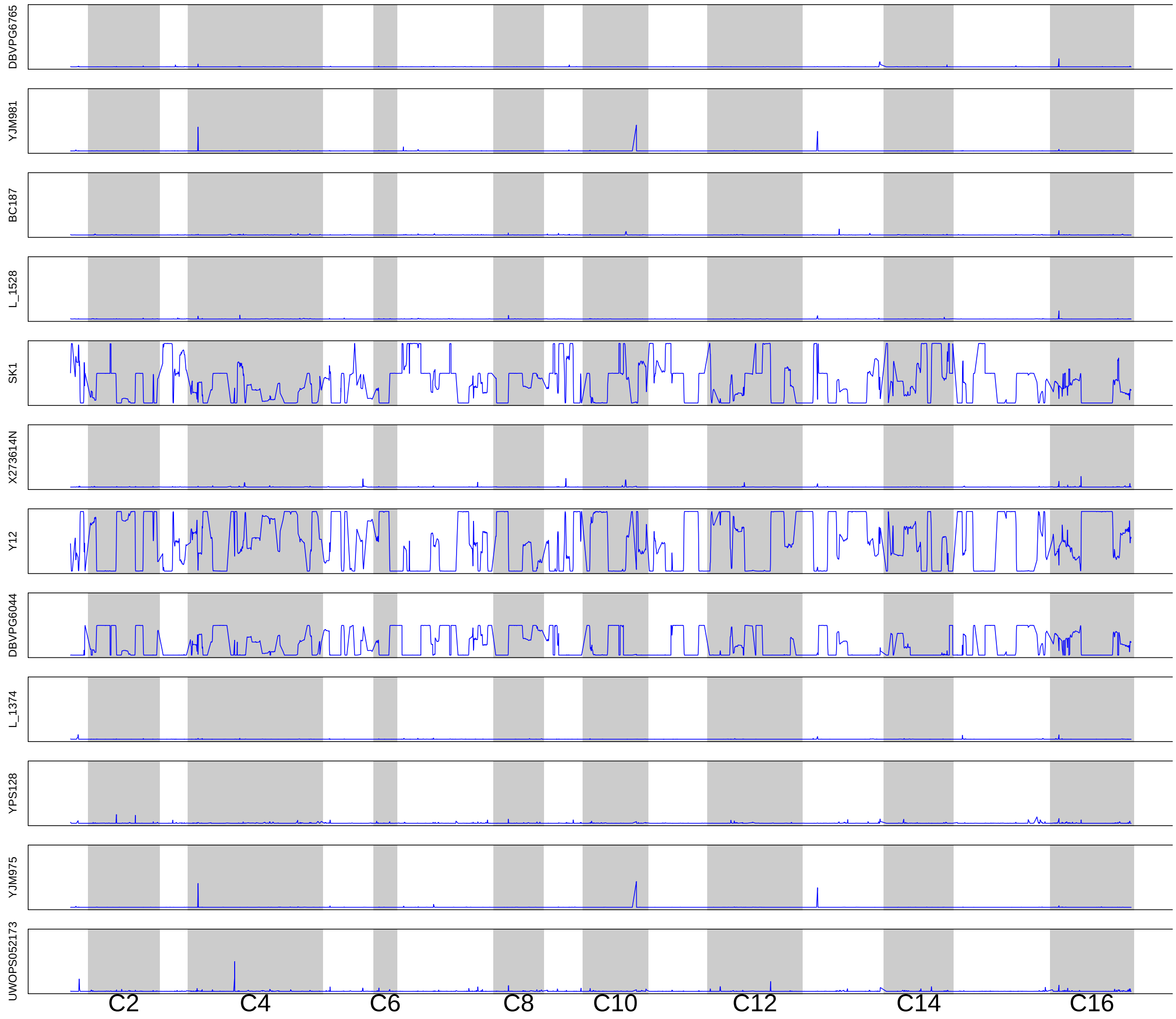

**Supplementary Figure S10.** Haplotype frequency estimates for population K12 after 12 cycles of outcrossing. Haplotype frequencies were estimated using sliding windows with a length of 30KB and a 1KB step size.

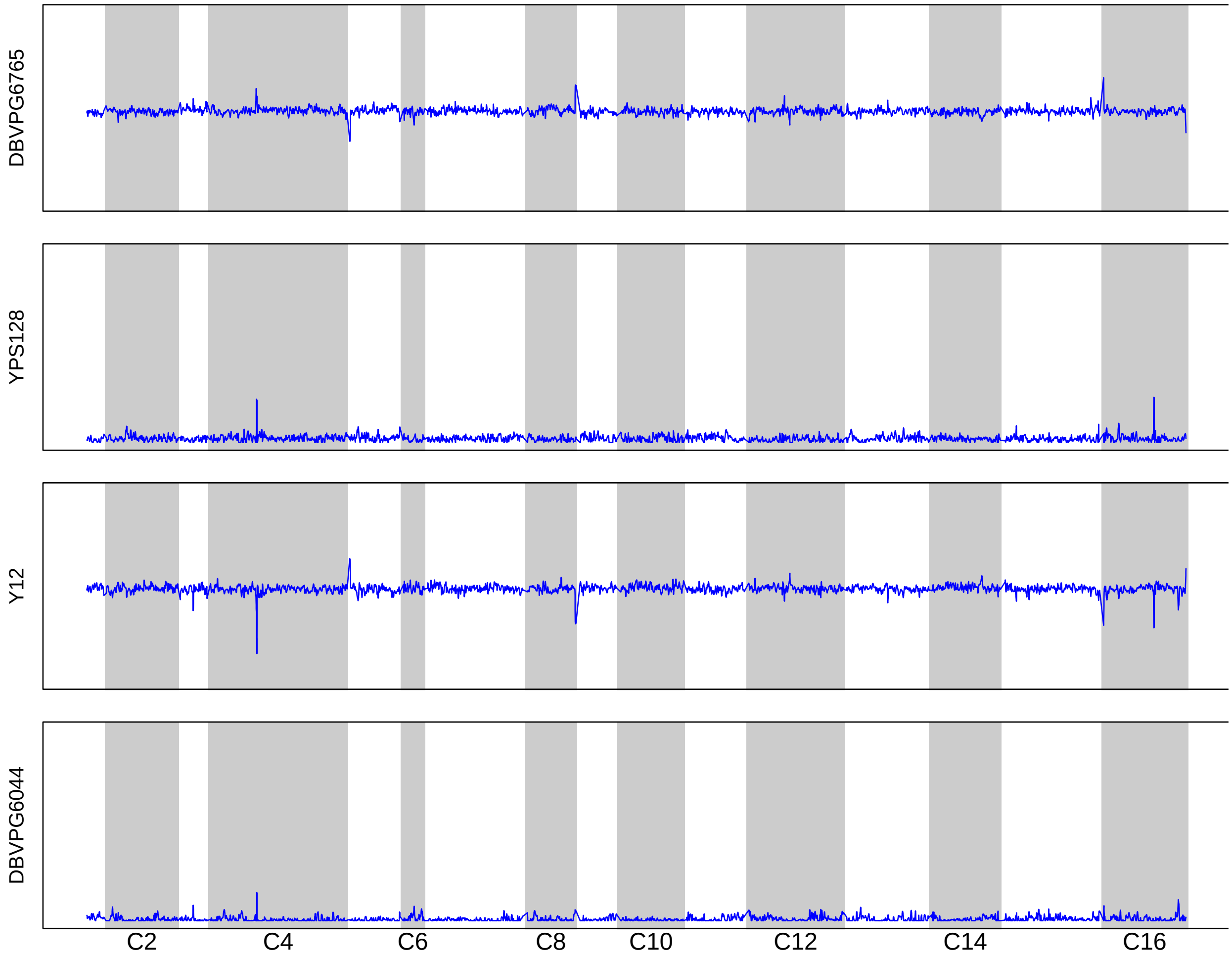

**Supplementary Figure S11.** Initial haplotype frequency estimates for population K4. Frequencies were estimated using sliding windows with a length of 30KB and a 1KB step size.

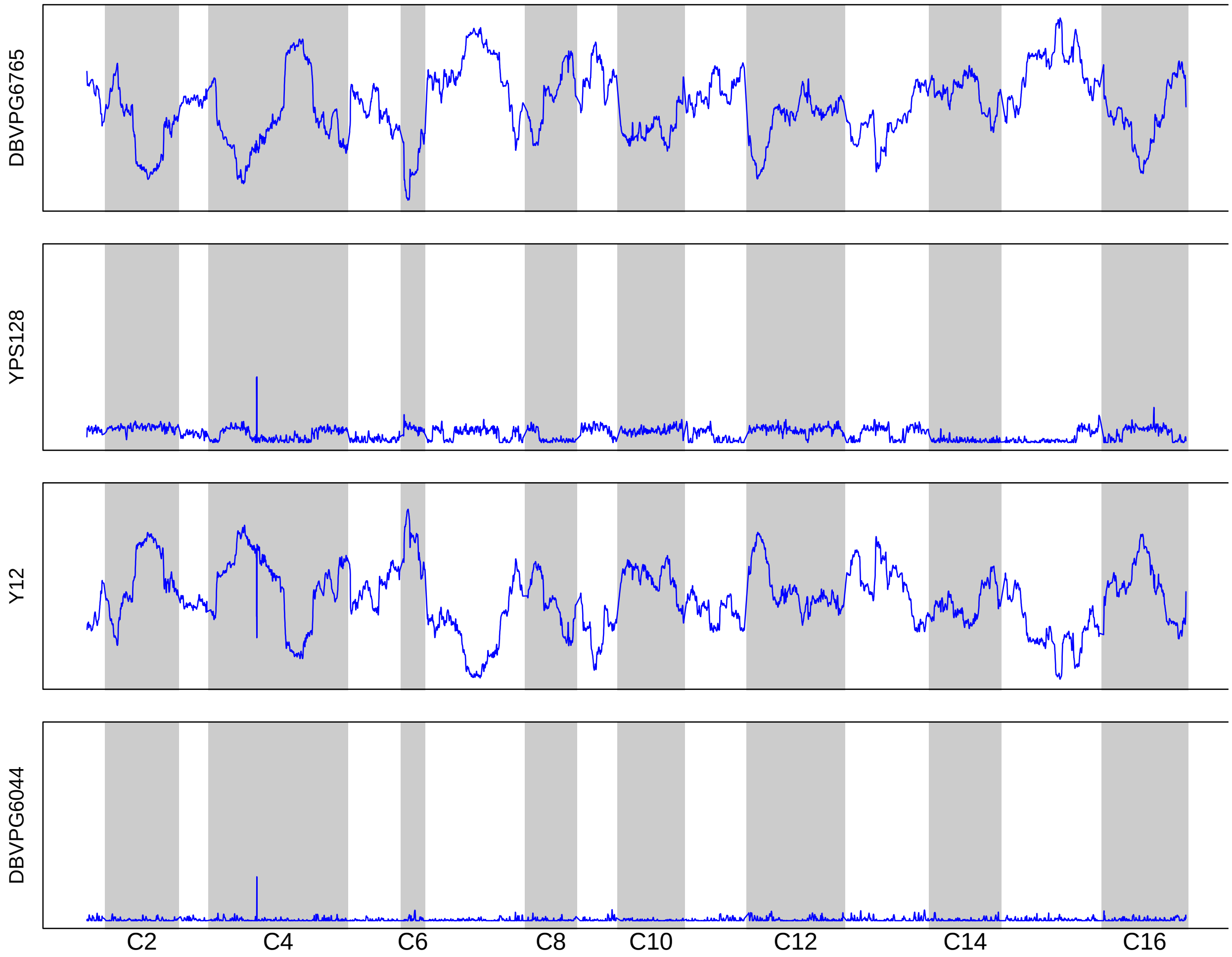

**Supplementary Figure S12.** Haplotype frequency estimates for population K4 after 6 cycles of outcrossing. Haplotype frequencies were estimated using sliding windows with a length of 30KB and a 1KB step size.

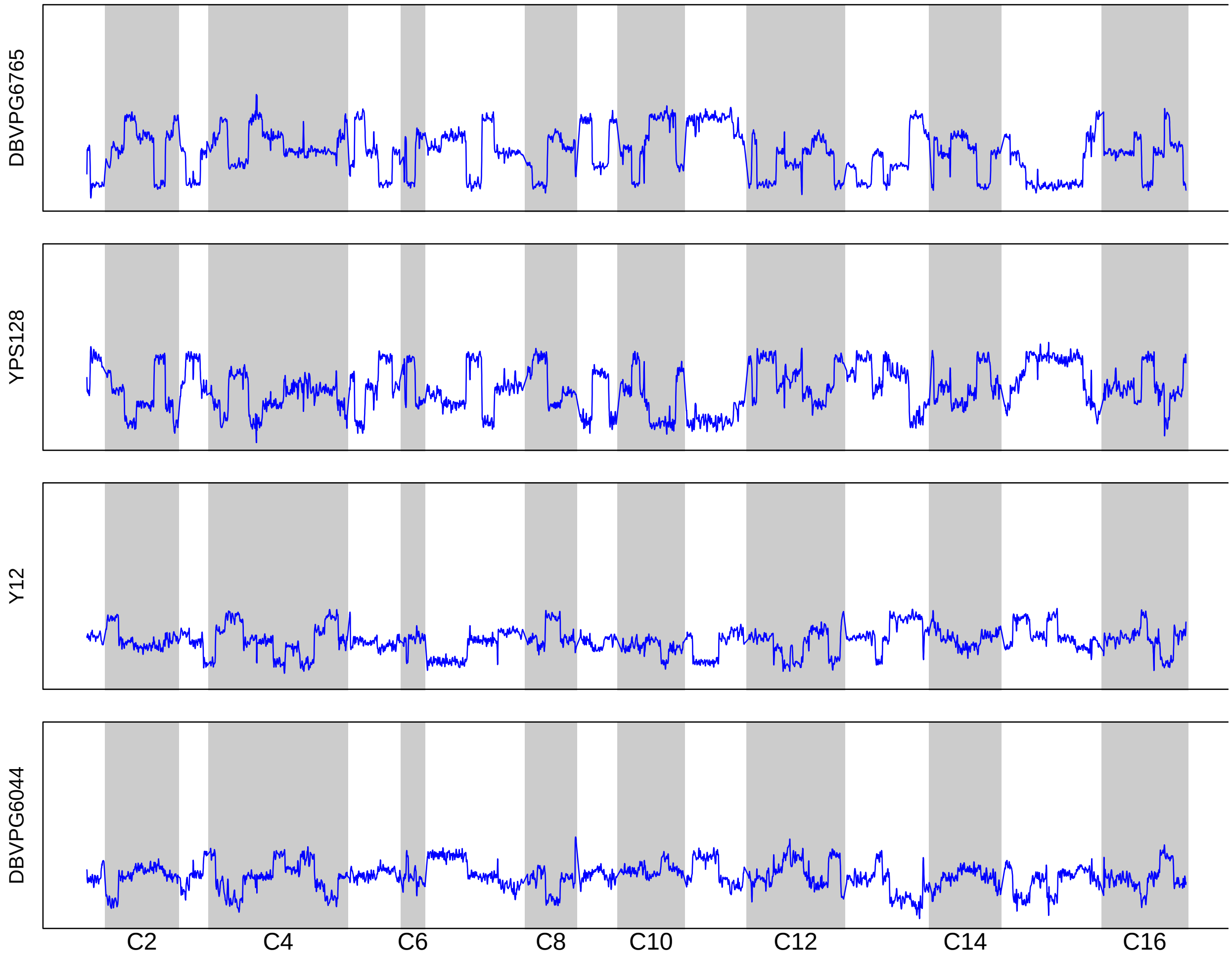

**Supplementary Figure S13.** Initial haplotype frequency estimates for population S4. Frequencies were estimated using sliding windows with a length of 30KB and a 1KB step size. Panels are ordered to match pairings in the first round of crossing (YPS128 was crossed with DBVPG6765, Y12 was crossed with DBVPG6044).

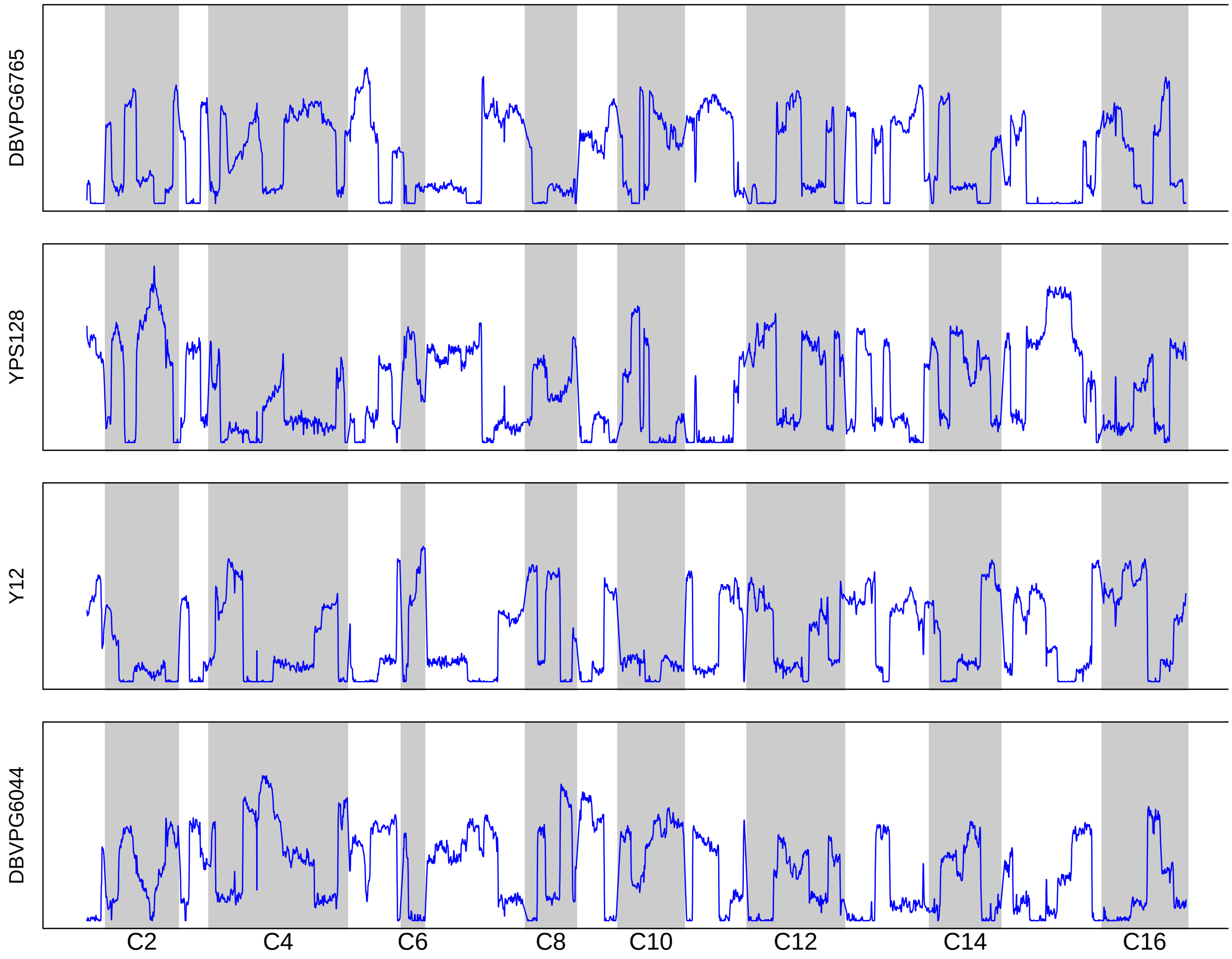

**Supplementary Figure S14.** Haplotype frequency estimates for population S4 after 6 cycles of outcrossing. Haplotype frequencies were estimated using sliding windows with a length of 30KB and a 1KB step size. Panels are ordered to match pairings in the first round of crossing (YPS128 was crossed with DBVPG6765, Y12 was crossed with DBVPG6044).

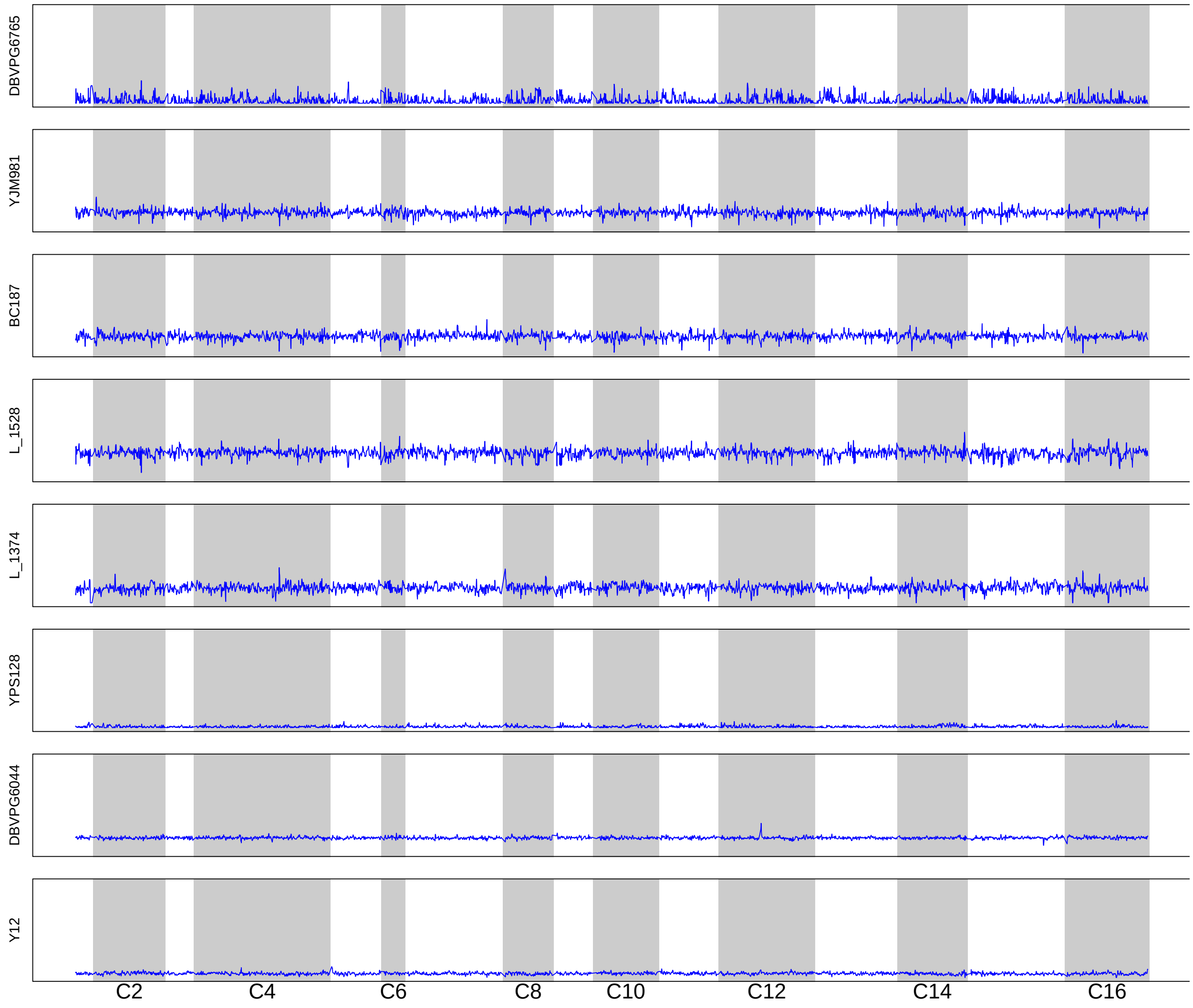

**Supplementary Figure S15.** Initial haplotype frequency estimates for population K8. Frequencies were estimated using sliding windows with a length of 30KB and a 1KB step size.

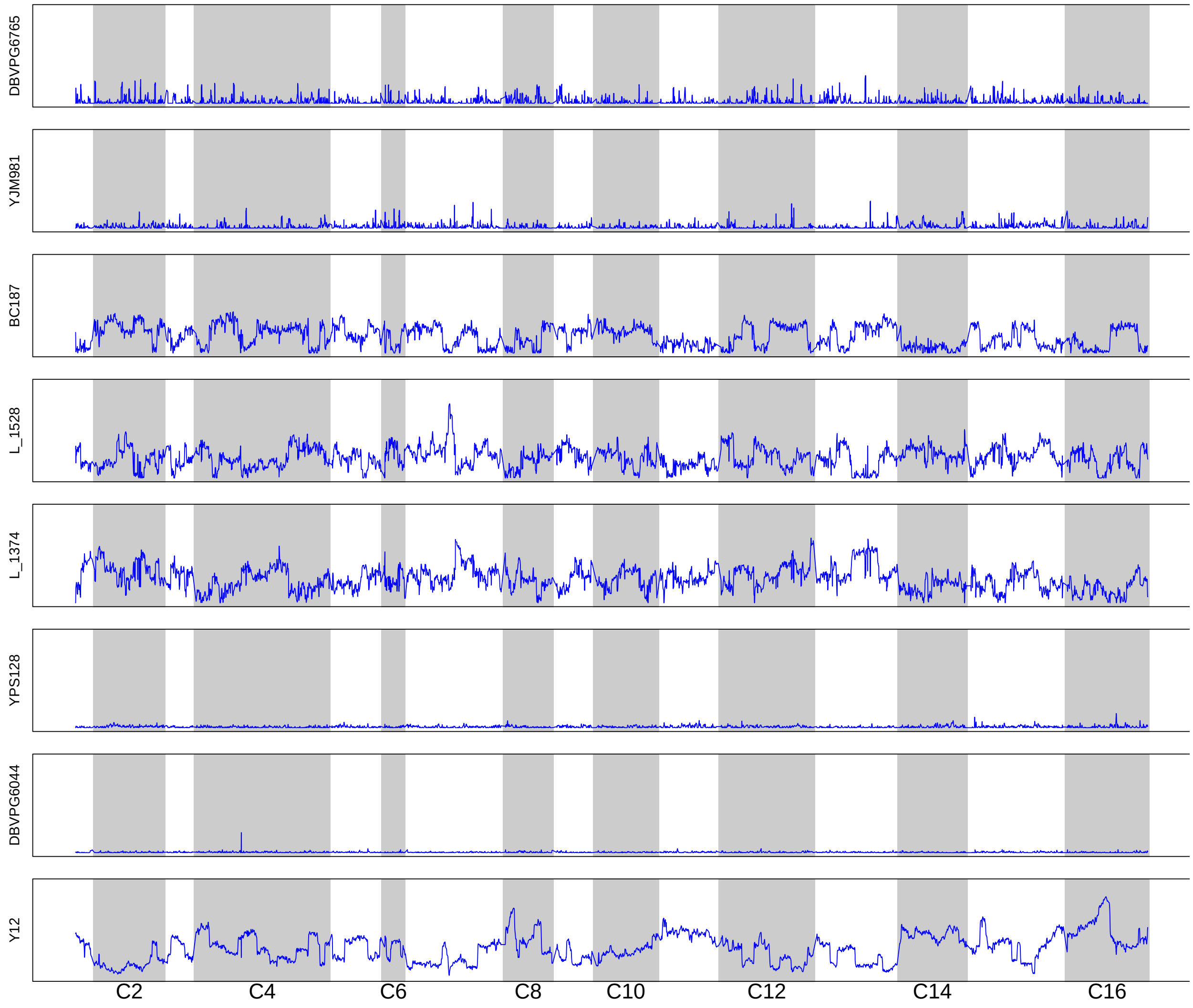

**Supplementary Figure S16.** Haplotype frequency estimates for population K8 after 6 cycles of outcrossing. Haplotype frequencies were estimated using sliding windows with a length of 30KB and a 1KB step size.

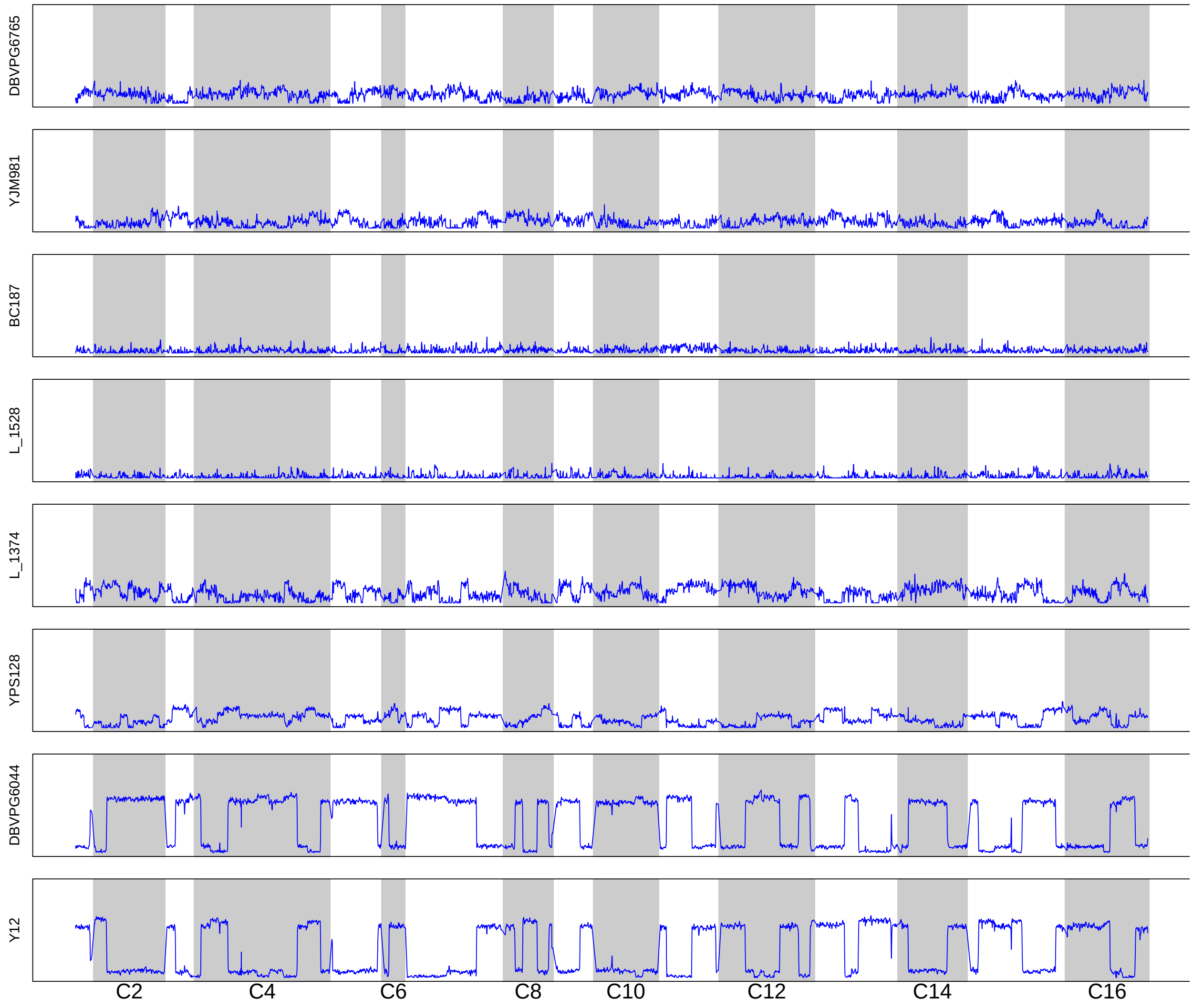

**Supplementary Figure S17**. Initial haplotype frequency estimates for population S8. Frequencies were estimated using sliding windows with a length of 30KB and a 1KB step size. Panels are ordered to match pairings in the first round of crossing (e.g. YJM981 was crossed with DBVPG6765, etc.)

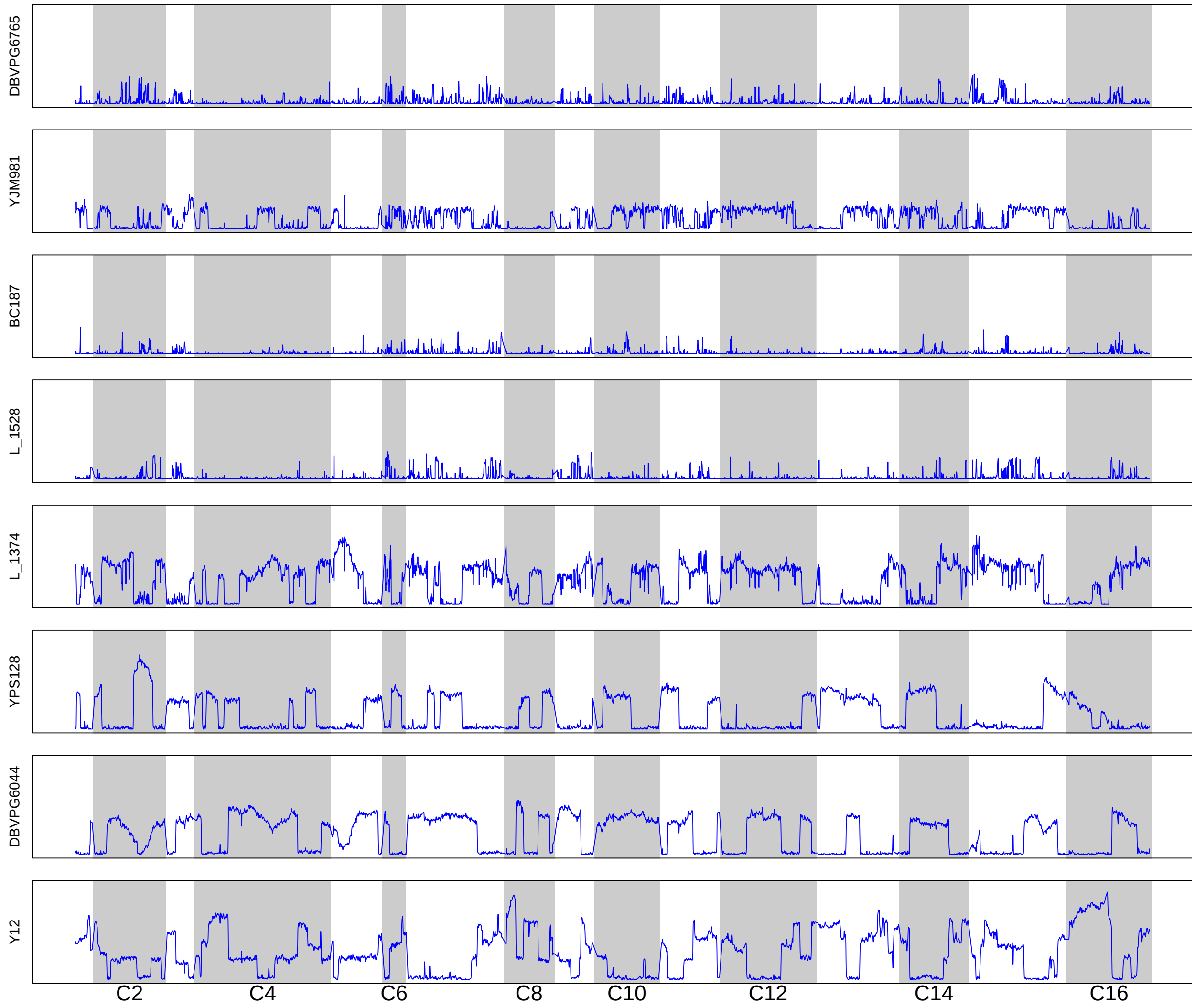

**Supplementary Figure S18.** Haplotype frequency estimates for population S8 after 6 cycles of outcrossing. Haplotype frequencies were estimated using sliding windows with a length of 30KB and a 1KB step size. Panels are ordered to match pairings in the first round of crossing (e.g. YJM981 was crossed with DBVPG6765, etc.).

### Supplementary Tables

**Supplementary Table S1.** Average genome-wide coverage at SNPs identified in each synthetic recombinant population. The total number of SNPs identified in each population (i.e. the number of positions at which read depth was averaged over) is shown in Table 1.

| **K-type populations** | | | **S-type populations** | | |
|---|---|---|---|---|---|
| Population | Outcrossing cycle | Mean Coverage | Population | Outcrossing cycle | Mean Coverage |
| K4 | 0 | 85 | S4 | 0 | 55 |
| K4 | 6 | 95 | S4 | 6 | 104 |
| K4 | 12 | 88 | S4 | 12 | 88 |
| K8 | 0 | 99 | S8 | 0 | 105 |
| K8 | 6 | 96 | S8 | 6 | 99 |
| K8 | 12 | 121 | S8 | 12 | 124 |
| K12 | 0 | 71 | S12 | 0 | 74 |
| K12 | 6 | 93 | S12 | 6 | 99 |
| K12 | 12 | 89 | S12 | 12 | 92 |

**Supplementary Table S2.** Mean genome-wide haplotype diversity for each synthetic recombinant population after 12 cycles of outcrossing. Variance in haplotype diversity and maximum expected values are also shown.

| **Population** | $\mathbf{H_{mean}}$ | $\mathbf{H_{var}}$ | $\mathbf{H_{max}}$ |
|---|---|---|---|
| S4 | 0.34 | 0.02 | 0.75 |
| K4 | 0.21 | 0.05 | |
| S8 | 0.63 | 0.02 | 0.86 |
| K8 | 0.65 | 0.01 | |
| S12 | 0.65 | 0.02 | 0.92 |
| K12 | 0.31 | 0.07 | |

**Supplementary Table S3.** Sporulation efficiencies in recombinant populations. Mean sporulation efficiencies were estimated as the proportion of cells that had formed tetrads within ~72 hours of incubation in minimal sporulation medium (~200 cells counted per replicate). Estimates are expressed as the mean of biological replicates (N=3 for the 4-founder populations; N=2 for the 8- and 12-founder populations), +/- standard deviation. Note that recombinant populations were assayed initially and after 12 cycles of outcrossing.

| Population | Sporulation efficiency (initial) | Sporulation efficiency (after 12 cycles of outcrossing) |
|---:|:---:|:---:|
| K4 | 0.35 +/- 0.09 | 0.43 +/- 0.22 |
| S4 | 0.40 +/- 0.15 | 0.33 +/- 0.17 |
| K8 | 0.40 +/- 0.14 | 0.55 +/- 0.12 |
| S8 | 0.35 +/- 0.05 | 0.42 +/- 0.04 |
| K12 | 0.29 +/- 0.01 | 0.10 +/- 0.04 |
| S12 | 0.36 +/- 0.08 | 0.30 +/- 0.12 |

**Supplementary Table S4.** Comparative growth rate estimates in recombinant populations and parental strains. Growth rate is expressed as mean doubling times in liquid YPD medium, which was averaged across 2 technical replicates and 2 biological replicates for all strains/populations of this study. Standard deviation of the biological replicates is reported alongside the mean. Note that recombinant populations were assayed initially and after 12 cycles of outcrossing. All strains/populations were assayed simultaneously.

| | **Doubling time (hrs) Initial** | **Doubling time (hrs) After 12 outcrossing cycles** |
|---|---|---|
| Diploid recombinant populations | | |
| K4 | 1.01 +/- 0.004 | 1.23 +/- 0.07 |
| S4 | 1.05 +/- 0.04 | 1.09 +/- 0.01 |
| K8 | 0.98 +/- 0.005 | 1.06 +/- 0.01 |
| S8 | 1.01 +/- 0.002 | 1.19 +/- 0.07 |
| K12 | 1.02 +/- 0.004 | 1.03 +/- 0.006 |
| S12 | 1.02 +/- 0.001 | 1.08 +/- 0.01 |
| Haploid founder strains | | |
| L_1528 | 1.01 +/- 0.01 | |
| UWOPS05_217_3 | 1.01 +/- 0.01 | |
| SK1 | 1.07 +/- 0.001 | |
| L_1374 | 1.08 +/- 0.01 | |
| YPS128 | 1.08 +/- 0.005 | |
| YJM981 | 1.11 +/- 0.09 | |
| DBVPG6044 | 1.15 +/- 0.01 | |
| BC187 | 1.15 +/- 0.01 | |
| DBVPG6765 | 1.19 +/- 0.001 | |
| Y12 | 1.21 +/- 0.06 | |
| 273614N | 1.22 +/- 0.01 | |
| YJM975 | 1.32 +/- 0.04 | |
